# Non-canonical regulation of glycogenolysis and the Warburg phenotype by soluble adenylyl cyclase

**DOI:** 10.1101/2020.02.02.929901

**Authors:** Jung-Chin Chang, Simei Go, Eduardo H. Gilglioni, Hang Lam Li, Hsu-Li Huang, Lonny R. Levin, Jochen Buck, Arthur J. Verhoeven, Ronald P.J. Oude Elferink

## Abstract

Cyclic AMP is produced in cells by two very different types of adenylyl cyclases: the canonical transmembrane adenylyl cyclases (tmACs, *ADCY1∼9*) and the evolutionarily more conserved soluble adenylyl cyclase (sAC, *ADCY10*). While the role and regulation of tmACs is well documented, much less is known of sAC in cellular metabolism. We demonstrate here that sAC is an acute regulator of glycolysis, oxidative phosphorylation and glycogen metabolism, tuning their relative bioenergetic contributions. Suppression of sAC activity leads to aerobic glycolysis, enhanced glycogenolysis, decreased oxidative phosphorylation, and an elevated cytosolic NADH/NAD^+^ ratio, resembling the Warburg phenotype. Importantly, we found that glycogen metabolism is regulated in opposite directions by cAMP depending on its location of synthesis and downstream effectors. While the canonical tmAC-cAMP-PKA axis promotes glycogenolysis, we identify a novel sAC-cAMP-Epac1 axis that suppresses glycogenolysis. These data suggest that sAC is an autonomous bioenergetic sensor that suppresses aerobic glycolysis and glycogenolysis when ATP levels suffice. When the ATP level falls, diminished sAC activity induces glycogenolysis and aerobic glycolysis to maintain energy homeostasis.

## Introduction

Since the discovery of 3’,5’-cyclic adenosine monophosphate (cAMP) by Earl Sutherland in his mechanistic study of glycogen metabolism (Sutherland & Rall, 1958), cAMP generated by transmembrane adenylyl cyclases (tmACs, ADCY1∼ADCY9) has been recognized as a versatile second messenger mediating many downstream signals of G-protein-coupled receptors (GPCRs). Recently, a new type of mammalian soluble adenylyl cyclase (sAC, *ADCY10*) was discovered, which is evolutionarily more conserved than tmACs and is localized not at the plasma membrane but intracellularly. (Chen, Cann et al., 2000, Kamenetsky, Middelhaufe et al., 2006). The intracellularly localized sAC and the plasma-membrane-localized tmACs share common cAMP effectors, such as protein kinase A (PKA) and “exchange factor directly activated by cAMP” (Epac1 and Epac2). However, sAC is not regulated by G-proteins and does not respond to forskolin (Kamenetsky et al., 2006). Instead, sAC activity is positively regulated by physiological concentrations of bicarbonate and fine-tuned by Ca^2+^ (Chen et al., 2000, Kleinboelting, Diaz et al., 2014, Litvin, Kamenetsky et al., 2003).

While the tmACs, regulated by G protein-coupled receptors in response to hormones and neurotransmitters, evolved subsequent to multicellularity, the evolutionary more ancient sAC-like enzymes first appeared in unicellular organisms and might regulate autonomous functions of cells. This concept is supported by the conservation of bicarbonate-mediated activation of sAC homologs in multicellular as well as in unicellular organisms (Kobayashi, Buck et al., 2004). As bicarbonate and CO_2_ are the primary product of substrate oxidation by the TCA cycle, we hypothesized that sAC regulates the autonomous metabolism of cells and this is supported by several characteristics of the enzyme. Firstly, sAC can directly sense the cellular energetic state due to its high *K*_m_ for its substrate ATP (ranging from 1 to 10 mM) (Jaiswal & Conti, 2003, Litvin et al., 2003, Zippin, Chen et al., 2013). Secondly, albeit at low level, sAC is expressed in almost all tissues examined (Geng, Wang et al., 2005, Levin & Buck, 2015). Thirdly, Ca^2+^, another versatile and universal second messenger, stimulates sAC in synergy with bicarbonate (Geng et al., 2005, Jaiswal & Conti, 2003), allowing sAC to tune the cellular metabolism according to ongoing cellular signaling. In line with this reasoning, the ATP- and calcium-sensing properties of sAC have been demonstrated to facilitate diverse metabolic regulations, including the regulation of oxidative phosphorylation by sensing CO_2_ production from the TCA cycle (Acin-Perez, Salazar et al., 2009) and/or free Ca^2+^ concentrations in the mitochondrial matrix (Di Benedetto, Scalzotto et al., 2013), induction of ATP-dependent insulin secretion in the pancreas (Zippin et al., 2013), secretion of aldosterone in adrenal cortex carcinoma H295R (Katona, Rajki et al., 2015), and TNF-induced respiratory burst in human neutrophils (Han, Stessin et al., 2005).

Over the past decade, it has become widely accepted that cAMP signaling is compartmentalized into microdomains which allows this single messenger molecule to mediate disparate functions even within a single cell. (Kamenetsky et al., 2006, Lefkimmiatis & Zaccolo, 2014). We have therefore examined the role of sAC-derived cAMP in energy metabolism and found that sAC acts as an acute switch for aerobic glycolysis and glycogenolysis in maintaining autonomous energy metabolism of cells. In multiple tested cell lines, acute suppression of sAC induces aerobic glycolysis, increased glycogenolysis and increased cytosolic NADH/NAD^+^ ratios. Apart from the established control of oxidative phosphorylation by sAC through regulation of complex I activity, we identified a novel regulatory mechanism of sAC on glycogen metabolism that is opposite to the well-established pathway of glycogenolysis by tmACs. Thus, while the tmAC-PKA signaling axis promotes glycogenolysis, a novel sAC-Epac1 signaling axis inhibits glycogen breakdown.

## Results

### Suppression of soluble adenylyl cyclase stably reprograms cell metabolism towards aerobic glycolysis

To investigate the role of sAC in autonomous metabolism of cells, we examined whether sAC activity regulates glycolysis at steady state. As sAC protein expression is generally low, we first used the SV-40 large T antigen immortalized human intrahepatic cholangiocyte cell line H69 (hereafter H69 cholangiocytes) (Grubman, Perrone et al., 1994), whose high endogenous expression of sAC was previously validated pharmacologically and genetically (Chang, Go et al., 2016). Consistent with previous studies, two sAC-specific inhibitors KH7 and LRE1 (Ramos-Espiritu, Kleinboelting et al., 2016b) suppressed cAMP production in H69 cholangiocytes (Figure S1A and S1B). Addition of KH7 or LRE1 elicited a dose-dependent increase in glucose consumption in H69 cholangiocytes. The increased glucose consumption was accompanied by increased lactate secretion and reduced pyruvate secretion, which resulted in an elevation of the lactate-to-pyruvate ratio in the medium in cells treated with sAC inhibitors (Figure 1A and 1B). The increased lactate-to-pyruvate ratio in media mirrored a similar change in intracellular lactate-to-pyruvate ratio (Fig. S2A). Similar changes in glucose, lactate and pyruvate were observed in sAC-knockdown H69 cholangiocytes (Figure 1C and 1D). Next, we investigated the general aspect of this observation with the human hepatoma cell line HepG2 (Fig. 1E and S2B) as well as with primary mouse hepatocytes and three additional cell lines of different tissue origin (human colorectal adenocarcinoma Caco-2, Abelson mouse leukemia virus-transformed macrophage RAW264.7, and mouse melanoma cell line B16F10; Figure S1C-F). Inhibition of sAC increased lactate secretion, and reduced pyruvate secretion in all four cell lines as well as primary mouse hepatocytes. Although the baseline lactate-to-pyruvate ratios were different in each cell type, sAC inhibition always led to an elevated medium lactate-to-pyruvate ratio. These findings show that both pharmacological and genetic suppression of sAC activity promotes aerobic glycolysis under normoxic conditions, resembling the Warburg effect observed in many cancer cells and consistent with the previously observed tumor suppression effect of sAC (Ramos-Espiritu, Diaz et al., 2016a).

**Figure 1.**
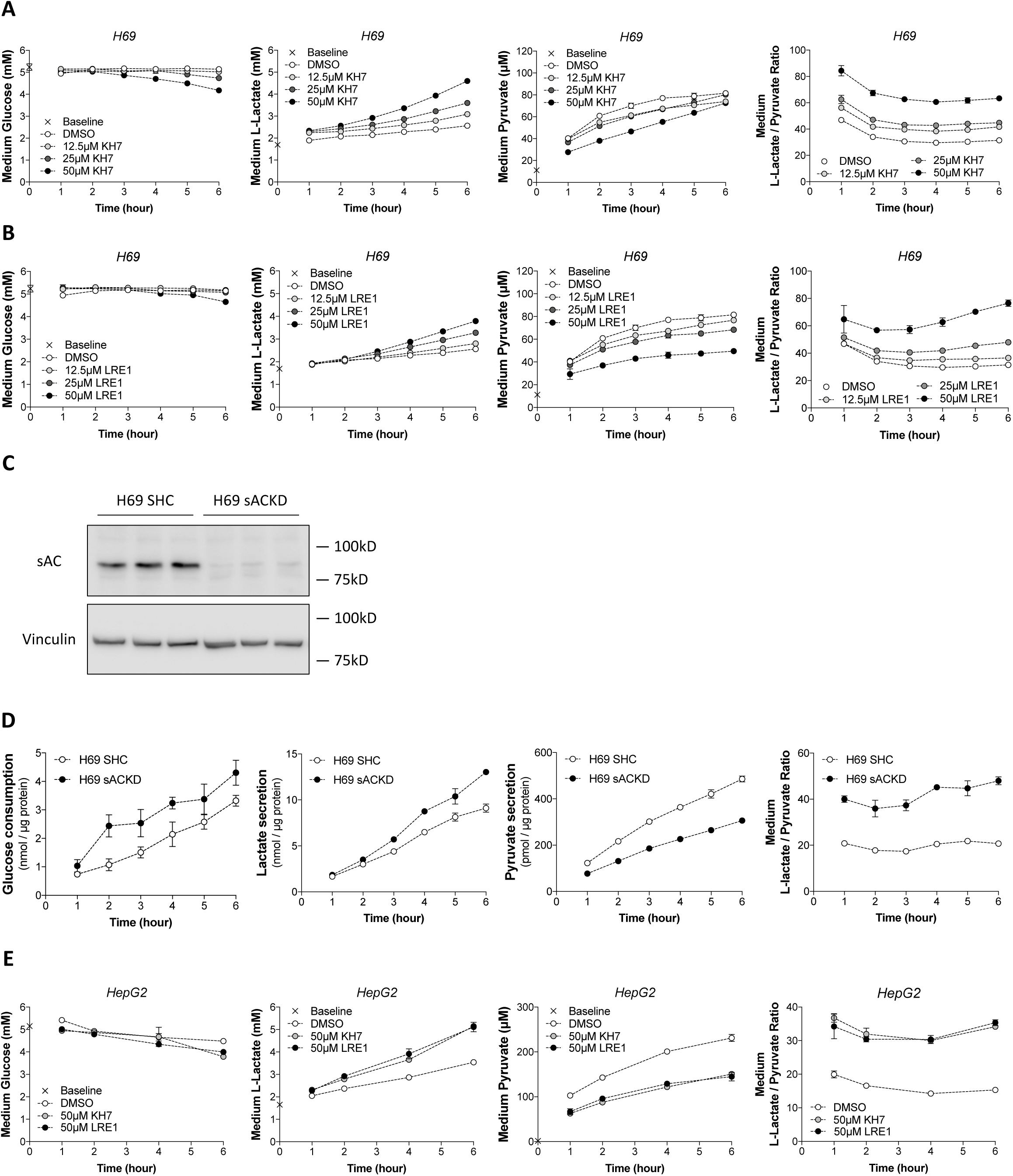
Suppression of soluble adenylyl cyclase stably reprograms cell metabolism towards a Warburg phenotype. (A-B) H69 human cholangiocytes were treated with different concentrations of sAC inhibitors KH7 (A) and LRE1 (B). Medium glucose, L-lactate, and pyruvate were sampled hourly for 6 hours and enzymatically determined. (C) Immunoblot of sAC in short hairpin control (SHC) and sAC-knockdown H69 human cholangiocytes. (D) Glucose consumption, L-lactate secretion, pyruvate secretion, and medium L-lactate-to-pyruvate ratio of SHC and sAC knockdown H69 human cholangiocytes. Results were normalized to protein contents. (E) HepG2 cells were treated with sAC-specific inhibitors KH7 and LRE1. Medium glucose, L-lactate and pyruvate were sampled after 1, 2, 4, and 6 hours and enzymatically determined. Data presented as mean ± SD (n = 3 for A, B and D; n=2 for E). Baseline value was the metabolite concentration in the blank medium. Data are representative of at least two independent experiments.

### Soluble adenylyl cyclase regulates the cytosolic NADH/NAD^+^ redox state and aerobic glycolysis via complex I

The monocarboxylate transporters (MCTs) on the plasma membrane equilibrates the extracellular lactate-to-pyruvate ratio with the intracellular lactate-to-pyruvate ratio (Halestrap & Wilson, 2012), which is in near equilibrium with the cytosolic redox couple NADH/NAD^+^ via lactate dehydrogenase (LDH) (Figure 2A). Since sAC inhibition increased the lactate-to-pyruvate ratio in both media (Figure 1) and inside cells (Figure S2A and S2B), we tested if sAC indeed regulates the cytosolic NADH/NAD^+^ redox state. To this end, we expressed the Peredox NADH/NAD^+^ biosensor (Hung, Albeck et al., 2011) in HepG2 cells to detect the ratio of free NADH over free NAD^+^ in the cytosol. The Peredox biosensor responded to various ratios, but not quantities, of lactate and pyruvate in the medium, confirming that the medium lactate-to-pyruvate ratio is in equilibrium with the cytosolic NADH/NAD^+^ ratio (Figure 2B). We found that the addition of LRE1 acutely and stably increased cytosolic NADH/NAD^+^ ratio in the presence of glucose (Figure 2C). As expected, when cells were acutely starved (Figure 2D) or when they were fueled with the short chain fatty acid octanoate alone, which supports oxidative phosphorylation in the absence of glycolysis (Figure 2E), the baseline cytosolic NADH/NAD^+^ ratio was lower than in the presence of glucose. Interestingly, under these two conditions without extracellular glucose, sAC inhibition strongly increased the NADH/NAD^+^ ratio in a transient fashion, indicating a rapid generation and subsequent depletion of glycolytic intermediates. These sAC-dependent changes in NADH/NAD^+^ ratio are dependent on glycolytic flux. When acutely starved cells were treated with 2-deoxyglucose to block glycolysis, the cytosolic NADH/NAD^+^ ratio reduced dramatically and the effect of the sAC inhibitor LRE1 was abolished (Figure 2F). Similarly, inhibiting glyceraldehyde-3-phosphate dehydrogenase (GAPDH), the NADH-generation step in glycolysis, with iodoacetate also significantly lowered cytosolic NADH/NAD^+^ ratio and blocked the effect of sAC inhibition (Figure 2G). These data demonstrate that cytosolic NADH is maintained by glycolysis and that sAC inhibition promotes glycolytic flux to increase the cytosolic NADH/NAD^+^ ratio, which becomes transient in the absence of glucose.

**Figure 2.**
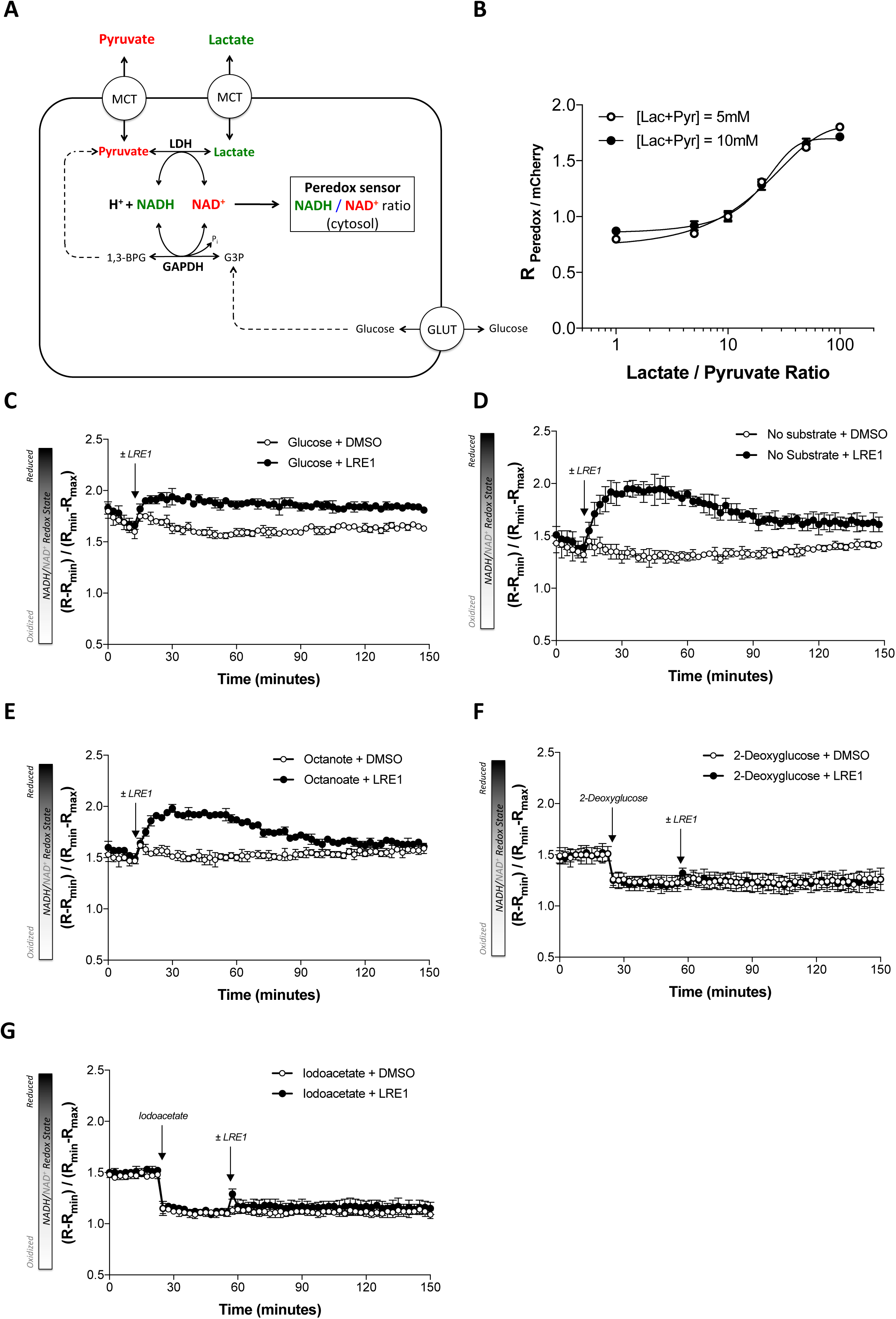
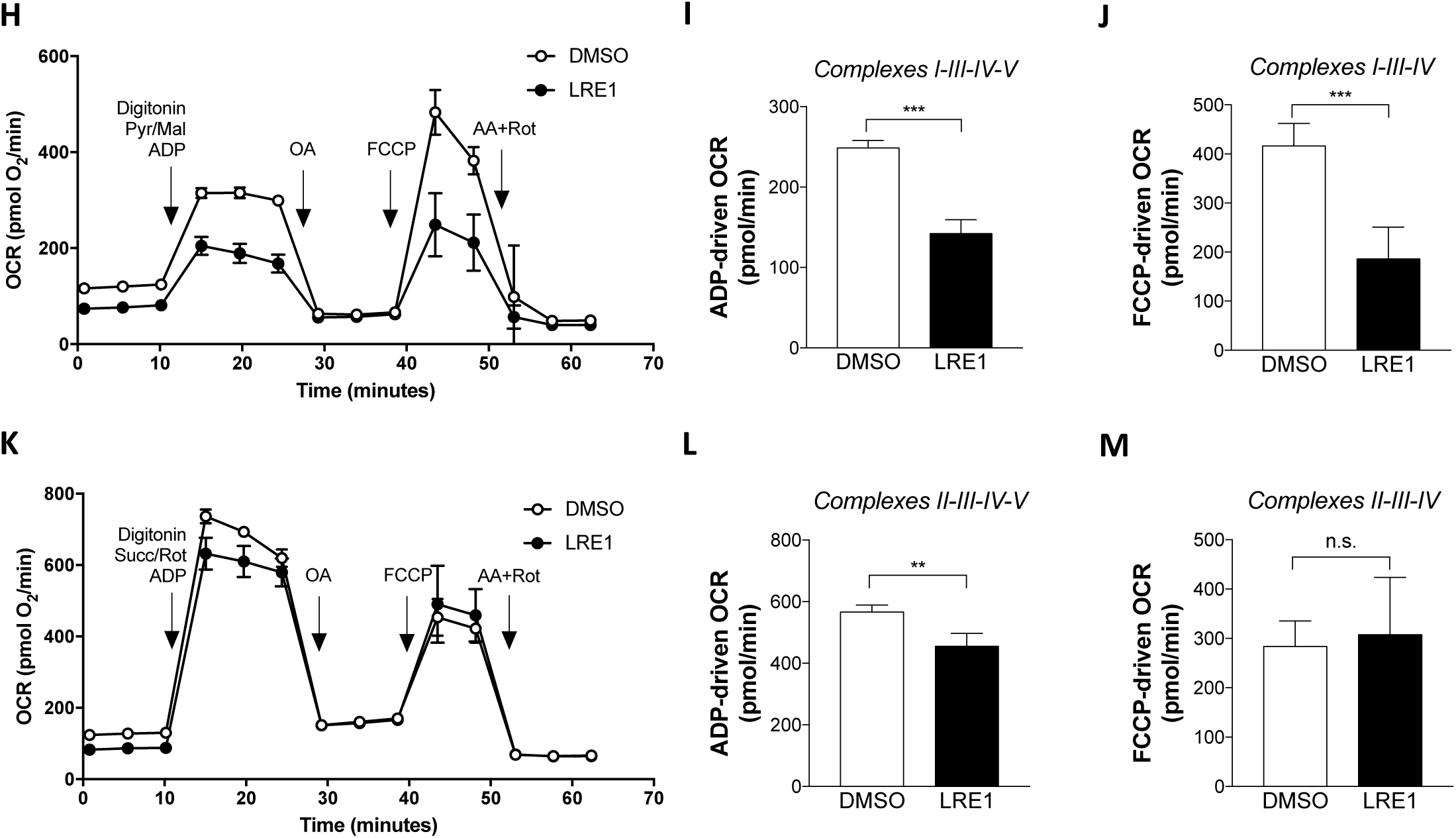
Soluble adenylyl cyclase regulates the cytosolic NADH/NAD^+^ redox state and aerobic glycolysis via complex I. (A) Ubiquitous monocarboxylate transporters (MCTs) equilibrate L-lactate and pyruvate across the plasma membrane. High lactate dehydrogenase activity in most cells brings the substrates (pyruvate and NADH) and products (lactate and NAD^+^) to near equilibrium. Therefore, the extracellular lactate-to-pyruvate ratio reflects the cytosolic lactate-to-pyruvate ratio and the cytosolic NADH/NAD^+^ ratio. (B) In *vivo* calibration of the Peredox-mCherry NADH/NAD^+^ redox sensor by media with different lactate-to-pyruvate ratio. (C-G) Cells were then pre-incubated for 1 hour with 5.5 mM glucose (C), no substrate (HBSS) (D, F, G) or 125 µM octanoate (E). Cells were refreshed again with the same medium and fluorescence ratio (R) of Peredox to tandem-tagged mCherry was monitored. For calibration, cells were incubated with 5 mM lactate and 5 mM pyruvate to obtain maximal (R_max_) and minimal (R_min_) ratios, respectively. The results were normalized to R_max_ and R_min_. Arrows indicate additions of 0.1% DMSO (vehicle control), 50 µM LRE1, 5.5 mM 2-deoxyglucose, or 0.5 mM iodoacetate. Results are presented as mean ± SD (n = 3-4). (H-M) HepG2 cells were pre-treated with 0.1% DMSO or 50 µM LRE1 for 10 minutes in HBSS and then the medium was changed to MAS buffer. Oxygen consumption rate (OCR) was monitored by Seahorse Flux Analyzer XF96. Cells were permeabilized by adding 25 µg/mL digitonin together with 1 mM ADP and substrates: (H-J) 2 mM pyruvate and 1 mM malate (n = 4 for DMSO and n = 5 for LRE1), and (K-M) 5 mM succinate and 2.5 µM rotenone (n = 4 for DMSO and n = 5 for LRE1). Subsequently, 2.5 µM OA, 1 µM FCCP, and 2.5 µM antimycin A (AA) + 2.5 µM rotenone (Rot) were injected as indicated. ADP-driven OCR (I and J) and FCCP-driven OCR (L and M) were derived from (H) and (K) as described in *Methods*. Data represent mean ± SD. Statistical analysis: (I-J, L-M) Two-tailed unpaired Student’s *t*-test. n.s., not significant, **P<0.01, ***P<0.001.

We next tested if sAC inhibition increased the cytosolic NADH/NAD^+^ ratio by decreasing NADH oxidation in mitochondria. While sAC has been reported to regulate various components of the mitochondrial respiratory chain in isolated mitochondria, the implicated target has not been consistent between the various reports (Acin-Perez et al., 2009, De Rasmo, Micelli et al., 2016, De Rasmo, Signorile et al., 2015). We reasoned that this might be due to differences in isolation procedures and the consequential disruption of mitochondrial networks. To minimize the disruption of the mitochondrial network and local structures, we examined the ADP-driven (state 3) respiration and FCCP-driven respiration (with ATP synthase blocked by oligomycin A) in digitonin-permeabilized HepG2 cells instead of isolated mitochondria. When mitochondria were fueled with complex I substrate (pyruvate), sAC inhibition significantly suppressed both ADP-driven and FCCP-driven respiration (Figure 2H-2J and S2C-S2E for pyruvate/malate and glutamate/malate, respectively). However, when mitochondria were fueled with complex II substrate (succinate), sAC inhibition only very mildly suppressed ADP-driven respiration and had no effect on FCCP-driven respiration (Figure 2K-2M). Thus, inhibiting sAC diminishes oxygen consumption primarily in the context of complex I driven respiration. In line with this, inhibiting complex I-dependent OCR with rotenone mimicked the Warburg-like phenotype observed in the sAC-suppressed state: increased lactate secretion, decreased pyruvate secretion, and elevated medium lactate-to-pyruvate ratio (Figure S2F-S2H). These results suggest that sAC regulates the cytosolic NADH/NAD^+^ redox state and aerobic glycolysis by affecting complex I activity.

### Soluble adenylyl cyclase is an acute switch for aerobic glycolysis that maintains energy homeostasis

Since sAC inhibition promotes glycolysis in intact cells and suppressed complex I activity in permeabilized cells, we asked if sAC can be an acute switch for aerobic glycolysis. To this end, we monitored glycolysis and oxidative phosphorylation by simultaneously measuring the extracellular acidification rate (ECAR) and oxygen consumption (OCR) in a Seahorse Flux Analyzer. As this method operates in ambient air and requires the use of nominally bicarbonate-free medium, we first examined whether endogenous bicarbonate production was sufficient to maintain sAC activity within the time frame of our experiments. We found that acute removal of CO_2_ in the gas phase did not affect sAC activity, suggesting sAC is primarily activated by endogenously produced bicarbonate (Figure S3A). In the presence of glucose, inhibiting sAC by LRE1 increased the extracellular acidification rate (ECAR) and decreased the coupled oxygen consumption rate (OCR) in both H69 (Figure 3A and 3B) and HepG2 cells (Figure 3C and 3D). Importantly, these metabolic effects of sAC inhibition took place within minutes and were stable. FCCP-uncoupled respiration was also suppressed upon sAC inhibition. To investigate whether energy homeostasis was maintained upon inhibition of sAC activity, we derived the ATP production rates of glycolysis and mitochondria from extracellular flux measurements in intact HepG2 cells following the method described by Mookerjee et al. (Mookerjee, Gerencser et al., 2017, Mookerjee, Goncalves et al., 2015). We found that in the presence of glucose, sAC inhibition acutely lowered ATP production by both oxidative phosphorylation and TCA cycle activity while increasing ATP production by glycolysis (Figure S3B-S3E). The overall ATP production remained unchanged (Figure 3SF). Direct determination of adenylate nucleotides in the cytosol also showed no changes in adenylate energy charge (Figure 3SG). These findings demonstrate that in the presence of glucose sAC maintains ATP homeostasis by reciprocally regulating ATP production via glycolysis and oxidative phosphorylation.

**Figure 3.**
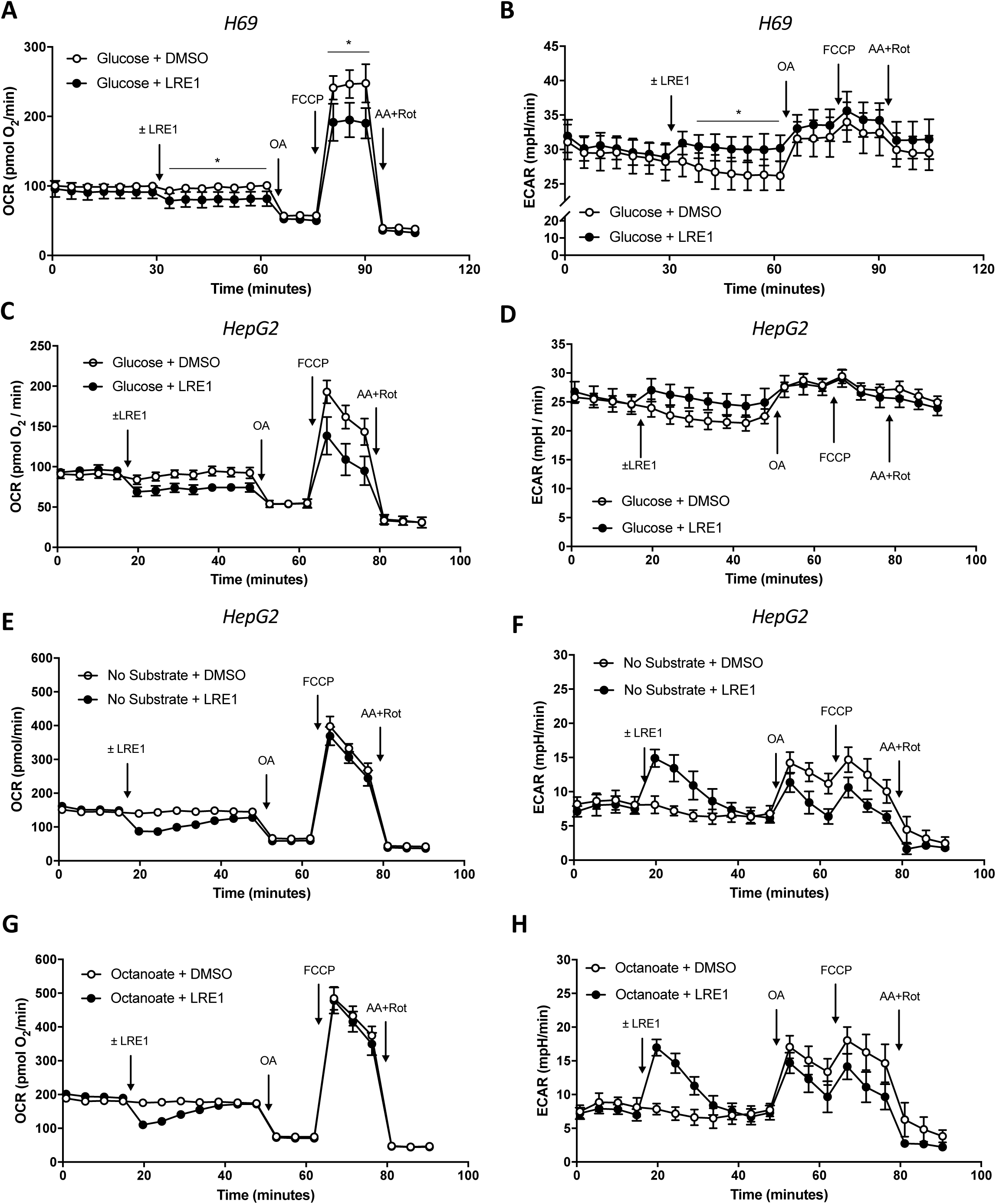
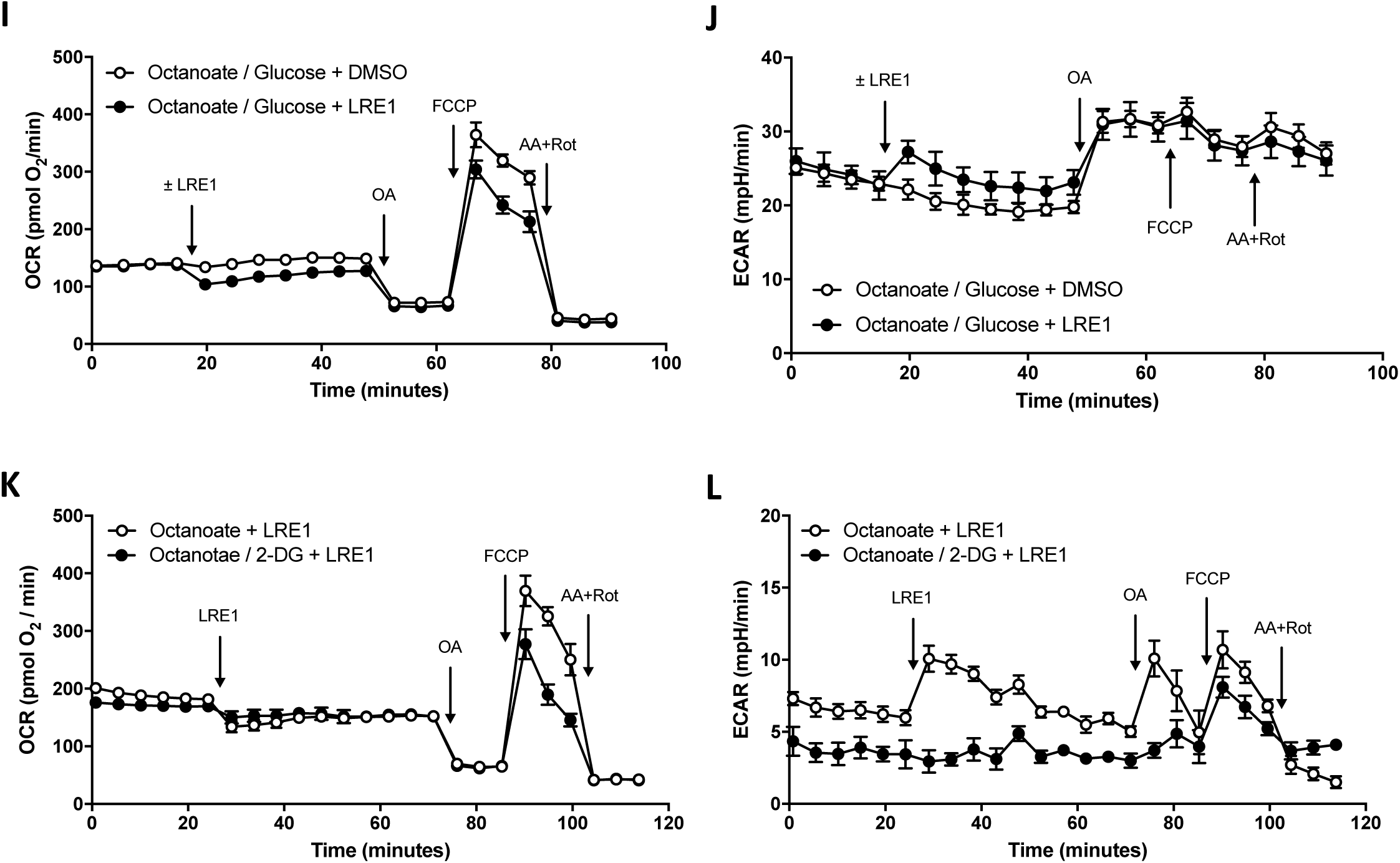
Soluble adenylyl cyclase is an acute switch for aerobic glycolysis that maintains energy homeostasis. The oxygen consumption rate (OCR) and extracellular acidification rate (ECAR) were analyzed by Seahorse Flux Analyzer XF96. (A-B) OCR and ECAR of confluent monolayers of H69 cholangiocytes were monitored in bicarbonate-free DMEM with 5.5 mM glucose. (C-D) HepG2 cells were pre-incubated for 60 minutes with HBSS with 5.5 mM glucose. OCR and ECAR were monitored. Arrows indicate the injection (in order) of 50 µM LRE1 or 0.1% DMSO, oligomycin A (OA), carbonyl cyanide-*p*-trifluoromethoxyphenyl hydrazone (FCCP), and antimycin A (AA) and rotenone (Rot). Data represent mean ± SD of (A-B) n = 5 and 5, (C-D) n = 10 and 9 for DMSO and LRE1, respectively. (E-L) HepG2 cells were preincubated for 1 hour in HBSS for ambient air without substrates (E-F), with 125 µM octanoate (G-H), with 5.5 mM glucose and 125 µM octanoate (I-J), and with 125 µM octanoate ± 5.5 mM 2-deoxyglucose (2-DG) (K-L). Oxygen consumption rate (OCR) and extracellular acidification rate (ECAR) were measured with Seahorse Flux Analyzer XF96. Arrows indicate the injection (in order) of 0.1% DMSO (vehicle control) or 50 µM LRE1, oligomycin A (OA), carbonyl cyanide-*p*-trifluoromethoxyphenyl hydrazone (FCCP), and antimycin A (AA) and rotenone (Rot). Data are presented as mean ± SD of (E-F) n = 10 and 8, (G-H) n = 10 and 11, (I-J) n=10 and 8 for DMSO and LRE1 respectively. Data in (K-L) represents mean ± SD of n = 8 and 6 for octanoate + LRE1 and octanoate/2-DG + LRE1 respectively. All data are representative of at least two independent experiments. Statistical analysis: (A-B) two-tailed unpaired Student’s *t*-test. *P<0.05.

We next examined how sAC regulates glycolysis and oxidative phosphorylation in the absence of glucose. When HepG2 cells were acutely starved (Figure 3E and 3F) or fueled solely with octanoate (Figure 3G and 3H), a membrane-permeant short-chain fatty acid and a direct substrate for mitochondrial β-oxidation, the effects of sAC inhibition on ECAR and coupled OCR remained reciprocal but became only transient. Calculation of the concomitant ATP production showed that ATP production by glycolysis was transiently stimulated upon sAC inhibition while ATP production by TCA cycle and oxidative phosphorylation were transiently suppressed (Figure S3H-S3K). Once again, as in the presence of glucose, the overall ATP production remained constant, resulting in an unchanged adenylate energy charge in the cytosol (Figure S3L and S3M). However, while sAC inhibition suppressed FCCP-uncoupled respiration in the presence of glucose, sAC inhibition did not affect FCCP-uncoupled respiration when cells were starved of glucose or fueled with octanoate. Thus, glucose is required for the stable inhibition of oxidative phosphorylation when sAC is inhibited under both coupled and uncoupled conditions. Indeed, when glucose was added to octanoate-fueled HepG2 cells, the effects of sAC inhibition on ECAR and OCR were stabilized (Figure 3I and 3J). Moreover, glucose also restored LRE1-induced suppression of FCCP-uncoupled respiration in octanoate-fueled HepG2 cells. Conversely, when HepG2 cells were fueled with octanoate in the presence of 2-deoxyglucose, the transient increase of ECAR was completely blocked and the degree of suppression of coupled OCR was reduced (Figure 3K and 3L). These data show that sAC inhibition suppresses oxidative phosphorylation by enhancing glycolysis in both the presence and absence of glucose, but in the absence of exogenously supplied glucose these changes are transient. The transient nature of these changes in the absence of glucose suggests that glycolytic intermediates may be responsible for the regulation of oxidative phosphorylation.

### Inhibition of soluble adenylyl cyclase promotes glycogenolysis

In the absence of exogenous glucose, ECAR was transiently stimulated by sAC inhibition, which implies that sAC inhibition increases availability of a glycolytic fuel. We hypothesized that these transient changes reflected the mobilization of a cellular glycolytic reserve, namely glycogen. While liver and muscles are the primary organs for glycogen storage, glycogen can be detected in tumor cell lines of various tissue origins (Rousset, Zweibaum et al., 1981). Indeed, we found that both HepG2 and H69 cells store significant quantities of glycogen (Figure 4A and 4B). In both cell lines, inhibition of sAC reduced glycogen stores within one hour. In HepG2 cells, sAC inhibition greatly reduced glycogen content in the absence of glucose but only mildly affected it in the presence of glucose (Figure 4A). In H69, sAC inhibition caused net glycogen breakdown both in the presence and absence of extracellular glucose (Figure 4B). Similarly, sAC inhibition promoted glycogen breakdown in acutely starved primary mouse hepatocytes, which could be prevented by inhibiting glycogen phosphorylase *a* with CP-91149 (Figure S4B). Furthermore, sAC inhibition also reduced glycogen content of primary mouse hepatocytes in the presence of glucose (Figure S4C). The induction of glycogenolysis in H69 human cholangiocytes and primary mouse hepatocytes upon sAC inhibition in the presence of glucose demonstrates that, at least in certain cell types, sAC-regulated glycogen turnover even plays a role in energy metabolism under normoglycemic conditions.

**Figure 4.**
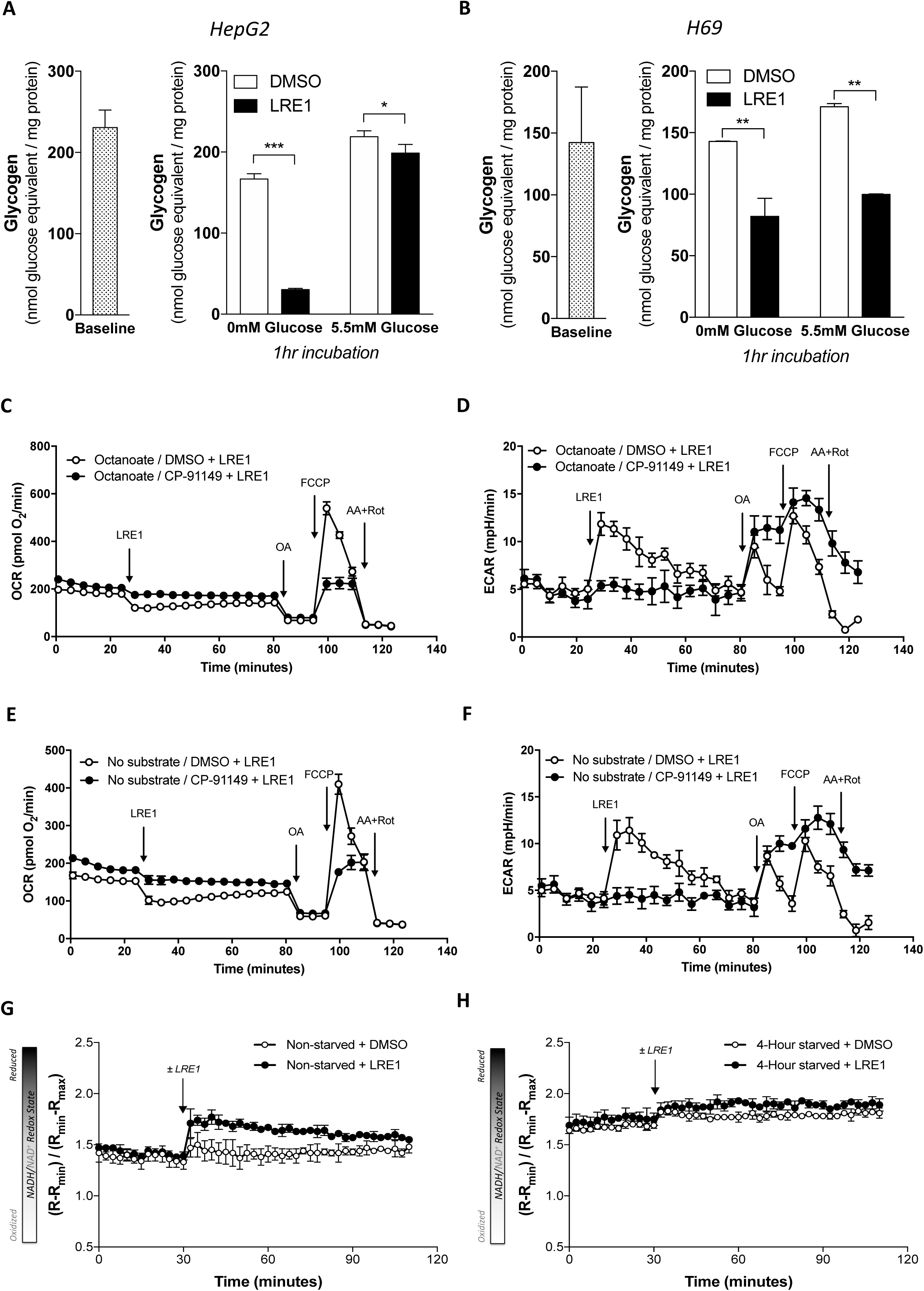
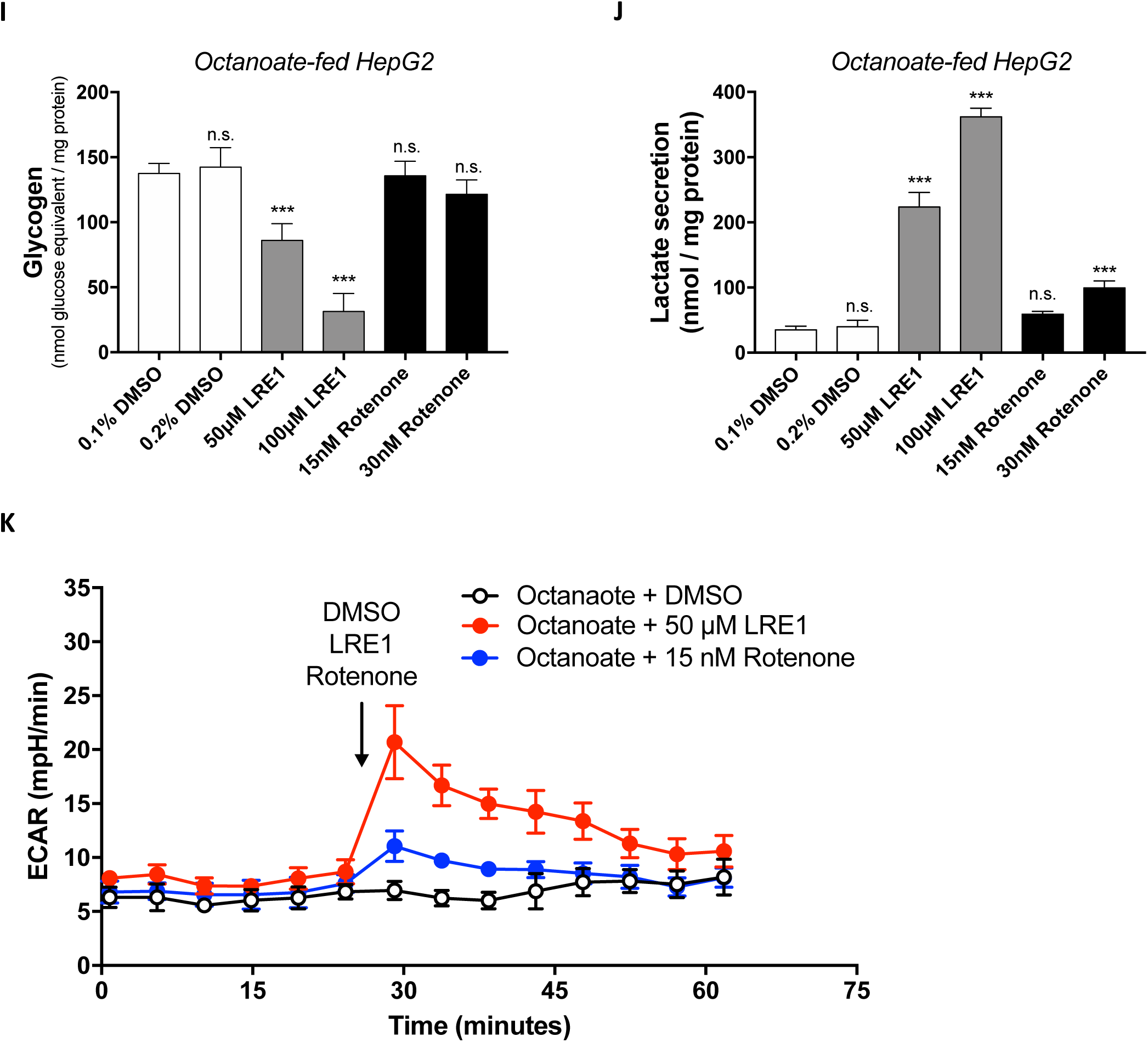
Inhibition of soluble adenylyl cyclase promotes glycogenolysis. (A) HepG2 cells were preincubated with 5.5 mM glucose for 1 hour. After harvesting baseline samples (n = 3), cells were refreshed with medium ± 5.5 mM glucose ± 50 µM LRE1 for another hour and then harvested for glycogen determination. Data represent mean ± SD (n = 3). (B) H69 cholangiocytes were treated as in (A) and glycogen content of cells were determined. Data presents mean ± SD of n = 8 for baseline and n = 2 for ± 5.5 mM glucose ± 50 µM LRE1. (C-D) HepG2 cells were pre-incubated with 125 µM octanoate in the presence or absence of CP-91149 for 1 hour. Oxygen consumption rate (OCR) and extracellular acidification rate (ECAR) were measured by Seahorse Flux Analyzer XF96. Arrows indicate the injection (in order) of 50 µM LRE1, oligomycin A (OA), carbonyl cyanide-*p*-trifluoromethoxyphenyl hydrazone (FCCP), and antimycin A (AA) and rotenone (Rot). Data are presented as mean ± SD of n = 5 and 5 for octanoate/DMSO and octanoate/CP-91149, respectively. (E-F) HepG2 cells were pre-incubated in HBSS without substrates in the presence or absence of CP-91149 for 1 hour. OCR and ECAR were measured as in (C) and (D). Data are presented as mean ± SD of n = 5 and 6 for no substrate/DMSO and no substrate/CP-91149, respectively. (G-H) Fluorescence ratio (R) of Peredox to tandem-tagged mCherry was monitored in non-starved or 4-hour starved HepG2 cells expressing Peredox-mCherry biosensor. Arrows indicate the addition of 0.1% DMSO (vehicle control) or 50 µM LRE1. For calibration, cells were incubated with 5 mM lactate and 5 mM pyruvate to obtain maximal (R_max_) and minimal (R_min_) ratios, respectively. The results were normalized to R_max_ and R_min_. (I-J) HepG2 cells were pre-incubated in glucose-free HBSS with 125 µM octanoate for 1 hour and then treated with 0.1% or 0.2% DMSO (vehicle control), 50 µM or 100 µM LRE1, and 15 nM or 30 nM rotenone for 15 minutes. Glycogen level (I) and secreted lactate (J) were assayed. Data are presented as mean ± SD (n = 3 for 0.1% DMSO condition of glycogen, n = 4 for the rest of the conditions) (K) HepG2 cells were pre-incubated with 125 µM octanoate 1 hour. Extracellular acidification rate (ECAR) was monitored by Seahorse Flux Analyzer XF96. The arrow indicates addition of 0.1% DMSO (vehicle control, n = 7), 50 µM LRE1 (n = 8), and 15 nM rotenone (n = 8). Data are presented as mean ± SD. Statistical analysis: (A-B) Two-way ANOVA with Tukey’s multiple comparisons test. (I-J) One-way ANOVA with Dunnett’s multiple comparisons test (against 0.1% DMSO). n.s., not significant, *P<0.05, **P<0.01, ***P<0.001.

We next examined whether glycogenolysis was directly responsible for the transient changes in ECAR, OCR, and cytosolic NADH/NAD^+^ ratio observed in the absence of extracellular glucose. In HepG2 cells fueled with only octanoate, the glycogen phosphorylase *a* inhibitor CP-91149 blocked the transient increase of ECAR evoked by sAC inhibition and significantly relieved suppression of OCR, showing that the transient increase of ECAR and suppression of OCR was the consequence of glycogenolysis (Figure 4C-4D). Similarly, in HepG2 cells acutely starved of substrates, CP-91149 also blocked the transient increase of ECAR by sAC inhibition, alleviated the transient OCR suppression, and preserved glycogen content (Figure 4E-4F and S4A). This confirms again that in the absence of exogenous glucose, LRE1-induced glycogenolysis fuels glycolysis, which in turn suppresses oxidative phosphorylation.

To test whether glycogenolysis-derived glycolysis was responsible for the transient increase in cytosolic NADH/NAD^+^ ratio upon sAC inhibition, we depleted glycogen by glucose starvation for 4 hours. In glycogen-depleted cells, sAC inhibition no longer increased the cytosolic NADH/NAD^+^ ratio (Figure 4G-4H, S4D). Thus, in the absence of glucose, sAC inhibition promotes glycogenolysis which increases glycolytic flux and the cytosolic NADH/NAD^+^ ratio.

Since sAC regulates complex I activity, we examined whether the induction of glycogenolysis upon sAC inhibition was merely the metabolic consequence of suppressed complex I activity or the result of parallel signaling events. To this end, we examined whether the complex I inhibitor rotenone could phenocopy the metabolic effect of sAC inhibition. In glucose-fed HepG2 cells, rotenone at 10-15 nM suppressed both coupled and uncoupled OCR to a comparable extent as 50 µM LRE1 (Figure S4E-G). However, rotenone only marginally induced glycogen breakdown and lactate secretion while LRE1 significantly depleted cellular glycogen and induced lactate secretion in octanoate-fed HepG2 cells (Figure 4I-J). Consistently, rotenone did not significantly increase ECAR in octanoate-fed HepG2 cells (Figure 4K). These data strongly suggest that sAC regulates glycogenolysis independently of its effect on complex I.

### sAC-cAMP-Epac1 and tmAC-cAMP-PKA form two distinct microdomains that signal opposite effects on glycogenolysis

The versatility of cAMP has led to the concept of signaling microdomains, where adenylyl cyclases, cAMP-degrading phosphodiesterases, cAMP effectors, and downstream substrates are brought together in one place to maintain specificity of cAMP signaling (Kamenetsky et al., 2006, Lefkimmiatis & Zaccolo, 2014). Our finding that sAC activity prevents rapid depletion of glycogen in the absence of glucose suggests that cAMP from sAC exerts an effect that is opposite to that of the well-established tmAC-cAMP-PKA axis. We confirmed these two opposing cAMP signaling pathways in HepG2 cells by demonstrating that both stimulation of tmACs by forskolin and inhibition of sAC by LRE1 promoted glycogenolysis (Figure 5A and 5B).

**Figure 5.**
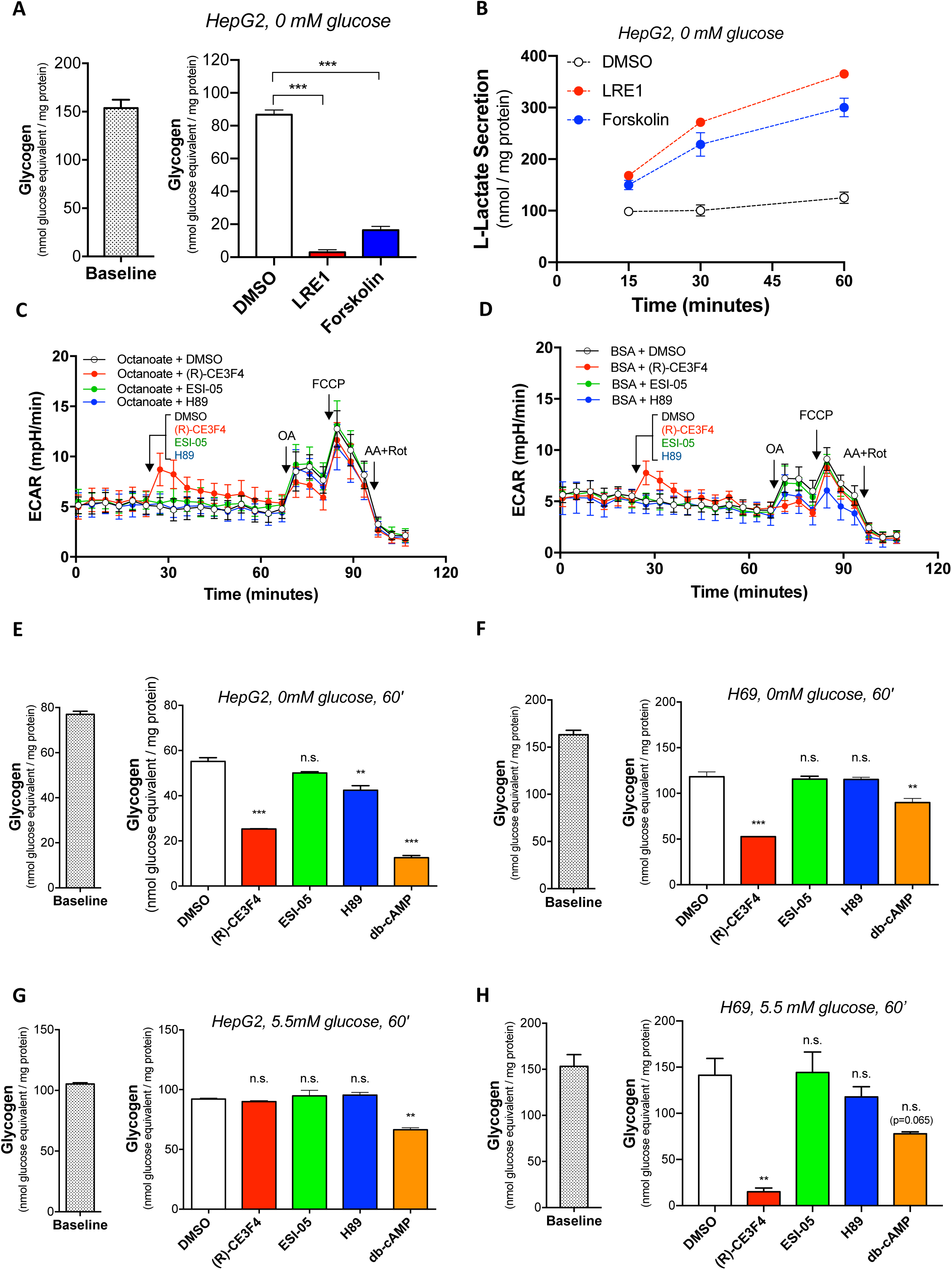
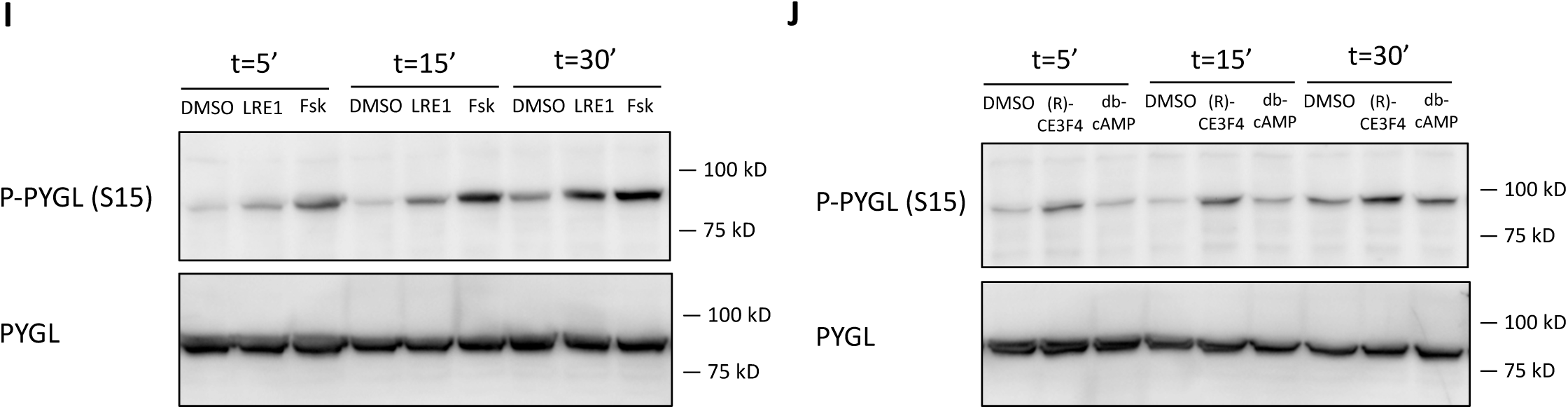
sAC-Epac1 and tmAC-PKA signaling have opposite effects on glycogen homeostasis. (A-B) HepG2 cells were acutely incubated in glucose-free HBSS with 0.1% DMSO (n = 3), 50 µM LRE1 (n = 3), and 1 µM forskolin (n = 3) for 60 minutes. Glycogen content at baseline (n = 6) and 60 minutes (A) and medium L-lactate at 15, 30, 60 minutes (B) were assayed. Data represent mean ± SD. (C) HepG2 cells were pre-incubated for 1 hour in HBSS for ambient air with 125 µM octanoate or 0.1% BSA. Extracellular acidification rate (ECAR) was measured with Seahorse Flux Analyzer XF96. After baseline measurement, 0.1% DMSO (vehicle control, n = 6), 50 µM (R)-CE3F4 (n = 6), 10 µM ESI-05 (n = 4), or 10 µM H89 (n = 5) were injected, followed by oligomycin A (OA), carbonyl cyanide-*p*-trifluoromethoxyphenyl hydrazone (FCCP), and antimycin A (AA) and rotenone (Rot). Data are presented as mean ± SD. (D) Same as (C) but in HBSS with 0.1% BSA. Data are presented as mean ± SD for 0.1% DMSO (vehicle control, n = 6), 50 µM (R)-CE3F4 (n = 6), 10 µM ESI-05 (n = 5), or 10 µM H89 (n = 5). (E-F) HepG2 cells (E) and H69 cholangiocytes (F) were pre-incubated in 5.5 mM glucose for 1 hour and acutely treated with 0.1% DMSO (vehicle control), 50 µM (R)-CE3F4, 10 µM ESI-05, 10 µM H89, or 100 µM dibutyryl-cAMP (db-cAMP) in glucose-free medium. Glycogen content was determined and normalized to total protein. Data are presented as mean ± SD (n = 2). (G-H) HepG2 cells (G) and H69 cholangiocytes (H) were pre-incubated in 5.5 mM glucose for 1 hour and acutely treated with 0.1% DMSO (vehicle control), 50 µM (R)-CE3F4, 10 µM ESI-05, 10 µM H89, or 100 µM db-cAMP in medium containing 5.5 mM glucose. Glycogen content was determined and normalized to total protein. Data are presented as mean ± SD (n = 2). (I) H69 cells were treated with 50 µM LRE1 or 1 µM forskolin for 5, 15, and 30 minutes. Phosphorylation of Ser 15 of liver form glycogen phosphorylase (PYGL) was examined by immunoblotting. (J) H69 cells were treated with 50 µM (R)-CE3F4 or 1 µM forskolin for 5, 15, and 30 minutes. Phosphorylation of Ser 15 of liver form glycogen phosphorylase (PYGL) was examined by immunoblotting. Statistical analysis: (A, E-H) One-way ANOVA with Dunnett’s multiple comparisons test (against DMSO group). n.s., not significant, **P<0.01, ***P<0.001.

We next investigated the mechanism underlying the disparity of cAMP effects derived from sAC and from tmACs on glycogen. sAC has been shown to signal via both Epac (Flacke, Flacke et al., 2013, Onodera, Nam et al., 2014) and PKA (Acin-Perez et al., 2009, Valsecchi, Konrad et al., 2017). Since tmAC-derived cAMP regulates glycogenolysis via PKA, we hypothesized that sAC-derived cAMP uses a different cAMP effector, namely Epac1 or Epac2. In glucose-starved HepG2 cells, i.e., fueled with only octanoate (Figure 5C) or acutely starved (Figure 5D), inhibition of Epac1 by the specific inhibitor (R)-CE3F4 (Courilleau, Bisserier et al., 2012) induced an acute, transient increase in ECAR with the same temporal and dynamic characteristics as sAC inhibition (Figure 3F and 3H), suggesting that Epac1 mediates sAC-dependent regulation of glycogenolysis. In contrast, the Epac2-specific inhibitor ESI-05 (Tsalkova, Mei et al., 2012) and the PKA inhibitor H89 did not induce significant ECAR changes.

To confirm the observed ECAR changes, we examined how Epac1, Epac2, and PKA regulate glycogenolysis by measuring the secretion of glycolytic end products and cellular glycogen levels. Indeed, in HepG2 cells acutely starved of glucose, only inhibition of Epac1 phenocopied the sAC-suppressed metabolic phenotype, which included increased lactate secretion, decreased pyruvate secretion, and increased medium lactate-to-pyruvate ratio (Figure S5A-S5C). Consistently, the Epac1-specific inhibitor (R)-CE3F4, but not the Epac2-specific inhibitor ESI-05 or the PKA inhibitor H89, induced significant glycogen breakdown in the absence of glucose in both HepG2 and H69 (Figure 5E and 5F, respectively). In addition, the PKA-selective activator dibutyryl-cAMP also promoted glycogen breakdown in both cell lines, confirming that PKA and Epac1 mediate opposite metabolic effects of tmAC and sAC signaling, respectively. Moreover, we observed that in the presence of glucose, the Epac1-specific inhibitor (R)-CE3F4 induced significant glycogen breakdown in H69 but not in HepG2 cells (Figure 5G and 5H), which mirrored the differential effects of sAC inhibition on glycogen in these cells (Figure 4A and 4B). Similarly, inhibition of sAC or Epac1 also induced glycogenolysis in several other cell lines (Figure S5G-S5I).

Glycogen content in cells is under the control of glycogen synthase and glycogen phosphorylase. Since inhibiting sAC-cAMP-Epac1 signaling promotes glycogenolysis in the absence of glucose, where glycogen synthase lacks substrates, we tested whether inhibition of sAC-cAMP-Epac1 signaling would increase glycogen phosphorylase activity. The activity of glycogen phosphorylase is regulated allosterically by AMP and covalently by phosphorylation of Ser15 by phosphorylase kinase. Since sAC inhibition did not cause changes in the adenylate energy charge in cytosol, we investigated if sAC-cAMP-Epac1 regulates glycogen phosphorylase by affecting its phosphorylation status at Ser15. We investigated this possibility in H69 cells because sAC-cAMP-Epac1 regulates the glycogen content of H69 cells also in the presence of a physiologic concentration of glucose. Indeed, both sAC-specific inhibitor LRE1 as well as tmAC-specific activator forskolin acutely increased Ser15 phosphorylation of glycogen phosphorylase (PYGL) (Figure 5I). Similarly, both the Epac1-selective inhibitor (R)-CE3F4 and the PKA-selective activator dibutyryl-cAMP induced acute phosphorylation of Ser15 in PYGL (Figure 5J). These data show that sAC-derived cAMP and tmAC-derived cAMP define independent signaling microdomains that have opposite effects on glycogenolysis: sAC-cAMP-Epac1 signaling suppresses glycogenolysis while tmAC-cAMP-PKA signaling promotes glycogenolysis.

## Discussion

Our present study demonstrates that the evolutionarily conserved sAC, which generates cAMP proportionately to intracellular levels of ATP, bicarbonate and free Ca^2+^, regulates the acute switch between aerobic glycolysis and glycogenolysis on the one hand and oxidative phosphorylation on the other (Figure 6). sAC-derived cAMP regulates complex I flux and ATP generation by oxidative phosphorylation; at the same time, and independently, sAC-derived cAMP signals via Epac1 to maintain glycogen levels. Via this metabolic tuning, sAC regulates the relative bioenergetic contributions of glycolysis, glycogenolysis, and oxidative phosphorylation to maintain a constant overall ATP production rate under different substrate availability and thereby maintains the adenylate energy charge.

**Figure 6.**
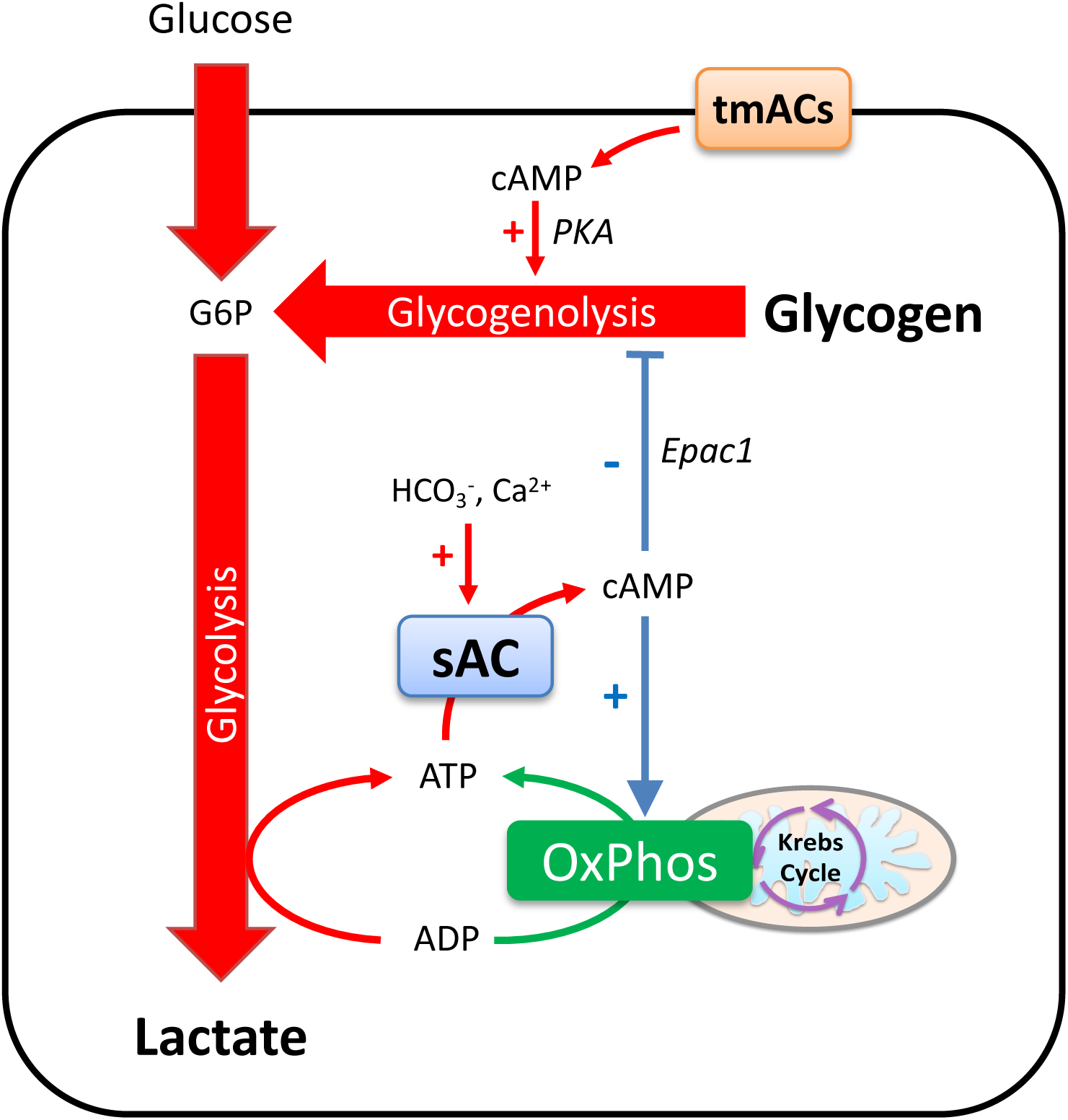
Working model of the acute regulation of aerobic glycolysis and glycogenolysis by soluble adenylyl cyclase. The activity of soluble adenylyl cyclase (sAC) is dependent on ATP and stimulated by Ca^2+^ and HCO_3_^-^. On the one hand, sAC regulates complex I activity; on the other hand, sAC-cAMP-Epac1 signaling suppresses glycogenolysis by reducing Ser15 phosphorylation of glycogen phosphorylase while the established tmAC-cAMP-PKA signaling activates glycogenolysis by increasing Ser15 phosphorylation of glycogen phosphorylase. When ATP level falls, decreased sAC activity induces a Warburg-like metabolic phenotype to maintain energy homeostasis, which is characterized by suppression of oxidative phosphorylation, promotion of glycolysis, and induction of glycogenolysis.

The primary metabolic signature of the sAC-suppressed status, as examined by pharmacological and genetic suppression of sAC, is a Warburg-like metabolic phenotype consisting of increased aerobic glycolysis, elevated cytosolic NADH/NAD^+^ ratio, and suppressed oxidative phosphorylation. In all cell lines tested and primary mouse hepatocytes, sAC inhibition acutely increased lactate secretion but decreased pyruvate secretion. The resulting increased lactate-to-pyruvate ratio in medium reflects an increase in the cytosolic NADH/NAD^+^ ratio, which was confirmed by experiments with the NADH/NAD^+^ biosensor Peredox. These data are consistent with the long-standing concept that the cytosolic NADH/NAD^+^ ratio can be inferred from the corresponding substrate-product pair of LDH (Veech, Raijman et al., 1970) and shows that sAC is a general regulator of the cytosolic NADH/NAD^+^ redox state. Most likely, sAC regulates the cytosolic NADH/NAD^+^ redox state by affecting complex I of electron transport chain, as evidenced by the results of our experiments with digitonin-permeabilized cells.

The second feature of the metabolic changes induced by sAC inhibition is the glucose dependency. In the presence of a physiologic concentration of glucose in the medium (5.5 mM), sAC inhibition simultaneously and stably increased the ECAR, elevated the cytosolic NADH/NAD^+^ ratio, and decreased the OCR. When glucose was removed from the medium, all the metabolic changes caused by sAC inhibition became transient, even if the cells were fueled with octanoate. Importantly, the transient changes in ECAR could be abolished by 2-DG and CP-91149, suggesting that the rise in NADH/NAD^+^ ratio by sAC inhibition is supported by glycogenolysis. Of note, the 2-DG and CP-91149 also greatly reduced the suppression of OCR by the sAC inhibitor, suggesting that the transient glycolytic flux from glycogenolysis also contributed to the suppression of oxidative phosphorylation. Importantly, these reciprocal changes in glycolysis and oxidative phosphorylation did not result in change of ATP production rate or the cytosolic adenylate energy charge, suggesting that sAC is a bioenergetic switch. In concordance with our data, Valsecchi *et al* reported that sAC knockout mouse embryonic fibroblasts also have increased ECAR and reduced oxidative phosphorylation (Valsecchi et al., 2017). They attributed this to reduced complex I activity in sAC knockout cells, while our data reveal that sAC regulates both complex I and glycogenolysis via independent mechanisms. In this way, sAC maintains bioenergetic homeostasis both in the presence of glucose and under acute glucose starvation.

The third and most important feature of the metabolic changes induced by sAC inhibition is a striking induction of glycogenolysis. Cellular glycogen levels are regulated by glycogen synthase and glycogen phosphorylase, both of which are regulated by phosphorylation as well as allosterically. Phosphorylation of glycogen synthase at serine residues 641, 645, and 649 by glycogen synthase kinase inactivates the enzyme (Roach, 1990). Glycogen phosphorylase exists in two states: the active R state and the much less active T state. Phosphorylation at serine 15 by phosphorylase kinase converts phosphorylase *b* to phosphorylase *a*, which has a favorable equilibrium for the active R state (Nolan, Novoa et al., 1964). These phosphorylation events are subject to simultaneous counter-regulation by protein phosphatase 1 (Cohen, 1983). In addition, glycogen phosphorylase *a* is allosterically inhibited by glucose while glycogen phosphorylase *b* is allosterically stimulated by AMP and inhibited by glucose-6-phosphate and ATP (reviewed in (Johnson, 1992)). On the other hand, glycogen synthase is allosterically activated by glucose-6-phosphate (Bouskila, Hunter et al., 2010). In the presence of glucose, high intracellular glucose and glucose-6-phosphate levels suppress stimulates glycogen synthase activity. Therefore, depending on glycogen synthase activity, the net result of suppression of sAC-cAMP-Epac1 in the presence of glucose may have variable effects on glycogen level in different types of cells. In the absence of glucose, glycogen synthase activity is reduced and inhibition of sAC-cAMP-Epac1 always results in rapid breakdown of glycogen, indicating increased glycogen phosphorylase activity. Our result show that inhibition of sAC-cAMP-Epac1 indeed promoted the conversion of phosphorylase *b* to phosphorylase *a*, which can be the result of either increased phosphorylase kinase activity or reduced protein phosphatase activity. Because sAC inhibition does not affect the adenylate energy charge of the cells, sAC-cAMP-Epac1 signaling is unlikely to allosterically activate glycogen phosphorylase by increasing AMP or decreasing ATP.

Our data demonstrate in cells of different tissue origins that cellular glycogen store is regulated by a novel sAC-cAMP-Epac1 signaling pathway that suppresses glycogenolysis. In contrast, the mobilization of glycogen store by epinephrine in muscle and by glucagon in liver both engage the classic tmAC-cAMP-PKA signaling. While the hormone-stimulated tmAC-cAMP-PKA mediates glycogenolysis in specialized tissues, the sAC-cAMP-Epac1 signaling seems to be more widely present and suppresses glycogenolysis and aerobic glycolysis to preserving glycogen store. From the view point of autonomous cellular metabolism, the sAC-cAMP-Epac1 would facilitate complete utilization of glycogen for ATP generation by both glycolysis and oxidative phosphorylation during glucose starvation.

With the discovery of multiple hormones that utilize the GPCR-G_αs_-tmAC-cAMP module for diverse cellular responses and the discovery of the intracellularly localized sAC, it is proposed that cAMP signaling is compartmentalized into microdomains to maintain specificity (Kamenetsky et al., 2006, Lefkimmiatis & Zaccolo, 2014). Cyclic AMP generated by sAC and by tmACs, albeit free to diffuse, is buffered by regulatory units of PKA(Agarwal, Clancy et al., 2016) and constantly degraded by phosphodiesterases (PDEs)(Jurevicius & Fischmeister, 1996, Zaccolo & Pozzan, 2002). The spatial specificity is further enhanced by A-kinase anchoring proteins (AKAPs), a family of diverse scaffolding proteins that assemble adenylyl cyclases, PDEs, PKA, protein phosphatase, and protein substrates on defined locations (Beene & Scott, 2007, Pidoux & Tasken, 2010). In addition, membranes impermeable to cAMP, such as the mitochondrial inner membrane(Di Benedetto et al., 2013), and lipid rafts in plasma membrane (Agarwal, Yang et al., 2014) also contribute to the spatial specificity. These mechanisms probably all contribute to the observed opposite effects of sAC and tmACs. For example, cAMP signaling by sAC promotes apoptosis (Chang et al., 2016, Kumar, Kostin et al., 2009) and compromises barrier function (Obiako, Calchary et al., 2013, Sayner, Alexeyev et al., 2006) while cAMP by tmACs protects against apoptosis (Chang et al., 2016, Kumar et al., 2009) and strengthens barrier function (Sayner et al., 2006). The differential signaling can also take place at the level of cAMP effectors. For instance, Epac1 activates Akt in a phosphatidylinositol 3-kinase-dependent manner while PKA suppresses Akt activation (Mei, Qiao et al., 2002). Our finding that sAC-cAMP-Epac1 has an opposing effect on glycogen metabolism as compared to tmAC-cAMP-PKA serves as yet another evidence for the cAMP microdomain signaling model.

Our findings also suggest that the metabolic regulation by sAC is potentially important in physiologic and pathologic processes associated with a switch of ATP production between glycolysis and oxidative phosphorylation, such as cancer metabolism (DeBerardinis & Chandel, 2016, Pavlova & Thompson, 2016), proliferation and differentiation of stem cells (Zhang, Nuebel et al., 2012), and activation of macrophages, dendritic cells and T-cells (Kelly & O’Neill, 2015). In particular, the metabolic reprogramming in the sAC-suppressed state, namely increased glycolytic flux and cytosolic NADH/NAD^+^ ratio with reduction in oxidative phosphorylation, resembles the seminal observation described by Otto Warburg that growing tumors utilized glucose by aerobic glycolysis despite sufficient oxygen tension for oxidative phosphorylation. This contention is supported by the observation that a large panel tumor samples have lower sAC expression than the corresponding normal tissues (Ramos-Espiritu et al., 2016a). Moreover, the recent development of NADH/NAD^+^ biosensors confirms that transformed cells have in general an elevated cytosolic free NADH/NAD^+^ ratio as compared to non-transformed cells (Schwartz, Passonneau et al., 1974, Zhao, Hu et al., 2015). While the constitutive Warburg effect in tumor cells is primarily driven by sustained genetic and epigenetic changes, we show here that sAC also acutely regulates energy metabolism and the Warburg-like metabolic phenotype. This mechanism may serve as a new therapeutic target to manipulate proliferation of cancer cells and inflammatory responses of macrophages, dendritic cells, and T-cells.

In summary, we demonstrate that the evolutionarily conserved soluble adenylyl cyclase is a master regulator of cellular energy metabolism, including the cytosolic NADH/NAD^+^ redox state, glycolysis, oxidative phosphorylation and glycogenolysis. It acts as a metabolic switch that may be explored for its therapeutic value in pathologic processes characterized by aberrant metabolic reprogramming.

## Acknowledgements

This study was supported by grant #11652-2018-1 from the Dutch Cancer Foundation (KWF/Alpe d’HuZes). The authors thank Dr. D. de Korte (Sanquin Blood Foundation, Amsterdam, NL) for performing the analysis of adenylate nucleotides. Dr. Buck and Levin are supported by NIH grants AG061290 and HD088571.

## Author Contributions

JCC, AJV and ROE developed the study concept. JCC, SG, EHG, HLL, HSH, and AJV designed, performed, and analyzed the experiments. JCC drafted the manuscript. All co-authors commented and revised the manuscript.

## Materials and Methods

### Materials

Unless otherwise indicated, materials were purchased from Sigma-Aldrich. Tet-pLKO-puro was a gift from Dmitri Wiederschain (Addgene plasmid #21915). pLKO.1-puro was a gift from Bob Weinberg (Addgene plasmid #8453). pCW-Cas9 was a gift from Eric Lander & David Sabatini (Addgene plasmid #50661). pcDNA3.1-Peredox-mCherry was a gift from Gary Yellen (Addgene plasmid #32383).

### Methods

#### Cell culture and lentiviral transduction

The immortalized human cholangiocytes H69 is kind gift from Douglas Jefferson (Grubman et al., 1994) and was cultured in DMEM/F-12 (3:1) with hormonal supplements as described (Chang et al., 2016). HepG2, Caco-2, B16F10, and Raw264.7 were cultured in DMEM (Invitrogen) with 2 g/L glucose, 1.8 g/L NaHCO_3_, 20 mM HEPES-NaOH pH 7.4 and 10% fetal bovine serum. For stable genetically modified cell lines, cells were transduced with lentivirus and selected with 2.5 µg/mL puromycin or 5 µg/mL basticidin. In acute transduction experiments, the transduced cells were used without puromycin selection. For all experiments, cells were refreshed the day before experiments.

#### Isolation and culture of primary mouse hepatocytes

Animal experiments were approved by the institutional animal experiment committee. Primary mouse hepatocytes were isolated from wild-type male C57BL/6 mice after overnight *ad libitum* feeding by a two-step collagenase perfusion method through the portal vein. Cells were cultured in collagen sandwich configuration overnight on a 60-rpm shaking platform in a 5% CO_2_, 37°C incubator. The isolation procedure and culture of primary mouse hepatocytes were performed as described (Gilglioni E.H., 2018).

#### Construction of lentiviral vectors and lentiviral production

The cloning procedure and short hairpin oligo sequences used to construct tetracycline-inducible lentiviral vector for sAC knockdown and scramble control has been described previously (Chang et al., 2016). Using pLKO.1-puro, the same procedure was followed to construct constitutively active lentiviral vectors for sAC knockdown and scramble control. To avoid formation of intracellular aggregation, we subcloned Peredox-mCherry NADH/NAD^+^ redox sensor for inducible expression. Briefly, puromycin resistance gene of pCW-Cas9 was removed by HincII and XbaI. Blasticidin resistance gene (Bsd) was amplified by forward primer 5’-CAGTCGGCTCCCTCGTTGACCGAATCACCGA CCTCTCTCCCCAGACGCGTATGGCCAAGCCTTTGTCTCA-3’ and reverse primer 5’-TTGTCCAGTCTAGACAT TGGACCAGGGTTTTCTTCAACATCACCACAAGTGAGGAGAGAACCTCTACCTTCATGCATGCCCTCCCA CACATAAC-3’. The purified PCR fragment was digested with HincII and Xbal and ligated with the open pCW-Cas9. The Cas9 insert was then removed by restriction enzymes NheI and BamHI. To create a multiple cloning site, sense oligo (5’-CTAGCGTCGACACCGGTGAATTCGTTTAAACCCTGCAGGG-3’) and antisense oligo (5’-GATCCCCTGCAGGGTTTAAACGAATTCACCGGTGTCGACG-3’) were annealed and ligated to the pCW backbone for pCW-MCS-BSD. The Peredox-mCherry insert was prepared by digesting pcDNA3.1-Peredox-mCherry with XbaI and PmeI. pCW-MCS-BSD was digested with NheI and PmeI and was ligated to the Peredox-mCherry insert for pCW-Peredox-mCherry-BSD. Lentivirus was produced in HEK293T using the third-generation packaging system.

#### Sample preparation and enzymatic assays for medium metabolites

Cells were refreshed with full culture medium one day prior to the experiment. On the day of the experiment, medium was changed to the experimental medium and cells were allowed to equilibrate in the experimental medium for one hour. The experimental medium was DMEM without glutamine, with 5.5 mM glucose and 10% (Figure 1) or 1% (Figure S1, Figure 5B) fetal bovine serum. In all other experiments, the experimental medium was based on modified Hank’s balanced salt solution (HBSS) for ambient air supplemented with 5.5 mM glucose and 0.1% fatty acid-free bovine serum albumin (BSA) (please refer to Suppl. Table 1 for formulation). We did not observe an effect of amino acids on the acute regulation of the Warburg-like phenotype and glycogen metabolism by sAC. At the start of the experiment, cells were exposed to media containing the indicated inhibitors and vehicle controls. At indicated time points, 50 µL spent medium was harvested and mixed with 75 µL ice-cold 5% (w/v) metaphosphoric acid (MPA) for deproteinization. After incubating for at least one hour at 4°C, the MPA-acidified samples were centrifuged at 20,000 × g for 10 minutes. The supernatants were harvested for enzymatic determination of L-lactate, pyruvate, and glucose. For the determination of intracellular lactate, pyruvate, and glucose-6-phosphate, medium was aspirated. Cells were washed once with ice-cold PBS and lysed in 3% ice-cold MPA. The acid homogenates were incubated at least one hour at 4°C and centrifuged at 20,000 × g for 10 minutes at 4°C. Lactate, pyruvate, and glucose are stable in 3% MPA at 4°C and were assayed directly from the MPA extracts.

#### Enzymatic assay for L-lactate, pyruvate, glucose

The enzymatic determinations of L-lactate, pyruvate and glucose were performed as previously described (Gilglioni E.H., 2018).

#### Sample preparation and enzymatic determination of glycogen

At the end of the incubation, cells were washed twice with ice-cold PBS and lysed in TTE buffer (1% Triton X-100, 10 mM Tris-HCl pH 8.0 and 1 mM EDTA-NaOH, pH 8.0) or RIPA buffer (150 mM NaCl, 20 mM Tris-HCl pH 8.0, 1% Triton X-100, 0.1% SDS, 0.5% Na-deoxycholate). Lysates were centrifuged at 20,000 × g at 4°C for 10 minutes. Supernatants were harvested for determination of glycogen and protein concentration. For the determination of glycogen, 50 µL of supernatant was mixed with 20 µL 0.35 M NaOH and heated at 80°C for 30 minutes to degrade monosaccharides. The hot alkali-treated lysates were then deproteinized by adding 30 µL 10% (w/v) MPA and incubated on ice for at least 1 hour or overnight at 4°C. Samples were centrifuged at 20,000 × g for 10 minutes. Twenty microliters of deproteinized samples or glucose standards (prepared in 3% MPA) were mixed with 100 µL solution A (0.75 mM homovanillic acid, 2.5 U/mL amyloglucosidase from *Aspergillus niger*, 50 mM K_2_HPO_4_-KH_2_PO_4_, pH 8.0). The resulting mixture has a pH of about 4.7, which is optimal for amyloglucosidase to hydrolyze glycogen. After 1-hour incubation at 45°C, 50 µL solution B was added (2 U/L horseradish peroxidase, 500 mM K_2_HPO_4_-KH_2_PO_4_, pH 8.0) to correct pH to 6.8 (assay condition for glucose). After determining the background fluorescence at λ_ex_/λ_em_= 320/450 nm in the CLARIOstar microplate reader (BMG LABTECH, Ortenberg, Germany), 50 µL start solution (8 U/mL glucose oxidase) was added and fluorescence was followed every 2 minutes until the reaction was completed (within 1 hour).

#### Determination of ATP, ADP, and AMP by high performance liquid chromatography (HPLC) for adenylate energy charges in cytosol

HepG2 cells were treated with 0.1% DMSO or 50 µM sAC-specific inhibitor LRE1 in HBSS containing indicated substrates. After indicated period of incubation, cytosolic metabolites were extracted with permeabilization buffer (120 mM KCl, 10 mM NaCl, 5 mM EDTA, 20 mM HEPES, pH 7.1) with 100 µg/mL digitonin. Two hundred microliter of digitonin extracts were mixed with 16 µL 70% perchloric acid for deproteinization and subsequently neutralized with 2.5 M K_2_CO_3_. The neutralized perchloric acid extracts of HepG2 cells were then analyzed for AMP, ADP and ATP by high-performance liquid chromatography using a Partisphere SAX column (Whatman International Ltd.) exactly as described (Bontekoe, van der Meer et al., 2017). Adenylate energy charge was defined as ([ATP] + 0.5 × [ADP]) / ([ATP] + [ADP] + [AMP]) according to Atkinson and Walton (Atkinson & Walton, 1967).

#### Cyclic AMP accumulation and ELISA

H69 cholangiocytes were cultured in 24-well plates. Cells were allowed to equilibrate with the indicated experimental medium for 1 hour and refreshed again with 200 µL experimental medium. Cells were pretreated with sAC-specific inhibitor KH7 or LRE1 (25 µL 10X solution) for 10 minutes and 500 µM 3-isobutyl-1-methylxanthine (25 µL 10X solution) was added to induce accumulation of cAMP. Accumulation was terminated by adding 100 µL 0.35 M HCl / 3.5% Triton X-100. The acidic homogenate was heated at 95°C for 10 minutes and centrifuged at 20,000 × g for 10 minutes at 4°C. The cAMP content in the supernatant was measured by the Direct cAMP ELISA kit (Enzo Life Sciences).

#### SDS-PAGE and Western blotting

Protein concentrations of whole cell lysates in RIPA buffer were quantified by BCA assay. Equal amount of protein (40-50 µg) was subjected to SDS-PAGE, transferred to PVDF membranes by semi-dry blotting and blocked overnight in 5% non-fat milk / PBST (phosphate-buffered saline with 0.05% (w/v) Tween 20) at 4°C. For immunodetection, the PVDF membranes were incubated with primary antibody for 1 hour, washed 3 times with TBST (Tris-buffered saline with 0.05% (w/v) Tween 20), incubated with horseradish peroxidase-conjugated secondary antibody for 1 hour, and washed again 4 times with TBST. All antibodies were diluted in 1% non-fat milk-TBST and incubation was performed at room temperature. The PVDF membrane was developed with homemade enhanced chemiluminescence reagents (100 mM Tris-HCl pH 8.5, 1.25 mM luminol, 0.2 mM p-coumarin and freshly added 3 mM H_2_O_2_) and detected using the ImageQuant LAS 4000 (GE Healthcare Life Sciences). Please refer to Table S2 for the list of primary and secondary antibodies and dilution.

#### Ratiometric fluorescence measurement of cytosolic NADH/NAD^+^ with Peredox-mCherry biosensor

Stable HepG2 cell lines harboring an inducible Peredox-mCherry construct or an empty vector (pCW-MCS-BSD) were cultured in clear-bottomed 96-well black plate (Costar 3603). Cells were induced with 0.8 µg/mL doxycycline for 48 hours before the experiment. Experiments were performed in HBSS modified for ambient air, supplemented with 0.1% BSA and indicated substrates at 37°C. For glycogen depletion, cells were starved in modified HBSS for 5% CO_2_ without glucose for 4 hours. Fluorescence of Peredox (F_1_) and mCherry (F_2_) was monitored at λ_Ex1_/λ_Em1_ = 400-20/535-70 (nm) and λ_Ex2_/λ_Em2_ = 570-20/645-80 (nm), respectively, in the CLARIOstar microplate reader. Responsiveness of the Peredox-mCherry sensor was confirmed in each experiment with a series of redox clamping solutions: 5 mM L-lactate, 5 mM pyruvate, and 5 mM L-lactate/pyruvate mixture at a ratio of 1, 5, 10, 20, 50, and 100. Signals from cells transduced with the empty vector (F_1_’ and F_2_’ for Peredox channel and mCherry channel, respectively) served to correct background. Background-corrected fluorescence ratio R was defined as (F_1_-F_1_’)/(F_2_-F_2_’). R was linearly transformed by setting the minimal fluorescence ratio R_min_ (5 mM pyruvate) and maximal fluorescence ratio R_max_ (5 mM L-lactate) to 1 and 2, respectively.

#### Extracellular flux analysis

Extracellular acidification rate (ECAR) and oxygen consumption rate (OCR) were measured with a Seahorse XF96 Analyzer (Agilent, United States). Cells were cultured in a 96-well Seahorse culture plate until confluence. Media were refreshed the day before the experiment. Cells were incubated 1 hour in experimental medium (modified HBSS for ambient air with 0.1% fatty acid-free BSA (see suppl. Table 1), with indicated substrates or inhibitors (2-deoxyglucose and glycogen phosphorylase *a* inhibitor CP-91149) in ambient air at 37°C and then refreshed with the same medium immediately before determination of ECAR and OCR. Twenty-five microliters of concentrated compound solutions prepared in experimental medium were injected as indicated. The final concentrations of compounds were as follows: LRE1, 50 µM; oligomycin A, 1.6 µM; FCCP, 1.1 µM; antimycin A, 4.62 µM; rotenone, 2.31 µM. The coupled respiration rate was defined as the difference between the average of the first three OCR measurements after the addition of inhibitors or vehicle control and the last OCR measurement after the addition of oligomycin A. The FCCP-driven respiration rate was defined as the difference between the first OCR measurement after FCCP addition and the last OCR measurement after the addition of oligomycin A.

#### Estimation of ATP production by glycolysis and by mitochondria using extracellular flux measurements

The time-lapsed ATP production rates were calculated from extracellular fluxes (ECARs and OCRs) as elegantly described by Mookerjee et al. (Mookerjee et al., 2017, Mookerjee et al., 2015). Since cells were pre-incubated in HBSS with glucose or octanoate alone until reaching steady state before imposing the acute perturbation of sAC activity, it was assumed that the oxidation of the provided substrates in the mitochondria was complete and that the contribution from other endogenous substrates (except for glycogen) was negligible. In cells fueled solely with octanoate, the glycolytic flux was assumed to come from glycogenolysis. The total ATP production rate (*J*_ATP_total_) is defined as the sum of the ATP production rate of glycolysis (*J*_ATP_glycolysis_, substrate level phosphorylation by phosphoglycerate kinase and pyruvate kinase) and the ATP production rate of mitochondria (*J*_ATP_mitochondria_, substrate level phosphorylation by succinyl-CoA synthetase in the TCA cycle and oxidative phosphorylation by the electron transport chain and ATP synthase). To calculate *J*_ATP_glycolysis_, it is necessary to distinguish the proton production by glycolysis from that of mitochondrial respiration. For this purpose, the total proton production rate at measurement time *t*, PPR_total_(*t*), is derived from ECAR by the following equation:

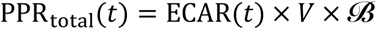

where ECAR(*t*) is the ECAR at measurement time *t*, *V* is the volume of the measurement micro-chamber (2 µL for Seahorse XF96 analyzer), and ***ℬ*** is the buffer capacity of the experimental medium. Since the experimental medium was buffered with 20mM HEPES-NaOH (pH 7.4), the contribution of other salt components and 0.1% BSA to buffer capacity were negligible. Taken the value of 7.31 as the p*K_a_* of HEPES at 37°C and an average assay pH of 7.35, the buffer capacity ***ℬ*** (M/pH) of the experimental medium was calculated to be 1.31×10^-2^ M/pH according to the definition of the buffer capacity (Urbansky & Schock, 2000):

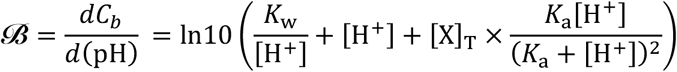

where *C*_b_ represents the normality of the added base, [X]_T_ represents the concentration of buffer, *K*_w_ represents ionization constant of H_2_O, and *K*_a_ represents the acid dissociation constant of the buffer.

The proton production rate of mitochondrial respiration (PPR_mito_) and glycolysis (PPR_gly_) at measurement time *t* is calculated as follows:

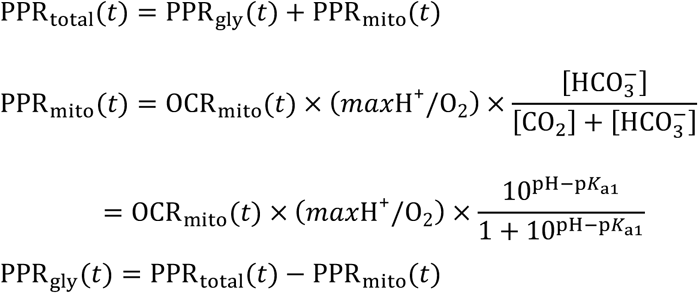

where *max* H^+^/O_2_ is the maximal proton production per O_2_ molecule consumed, and p*K*_a1_ is the combined equilibrium constant (6.093 at 37°C) for the hydration of CO_2_ and the dissociation of H_2_CO_3_. The OCR_mito_ (needed for the calculation of PPR_mito_) and OCR_coupled_ (needed for the calculation of *J*_ATP_mitochondria_) at measurement time *t* are derived from the measured OCR, OCR(*t*), as follows:

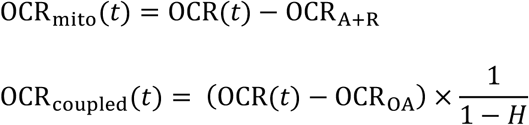

where OCR_OA_ and OCR_A+R_ are the last of the three OCR measurements after addition of oligomycin A and the combination of antimycin A and rotenone, respectively. Because oligomycin A increases the mitochondria membrane potential, proton leak and the associated oxygen consumption will increase. This will lead to an underestimation of the coupled OCR.

Therefore, a correction for hyperpolarisation is included, where *H* represents the percentage of underestimation and is taken to be 9.2% according to Affourtit and Brand (Affourtit & Brand, 2009). The OCR_coupled_ for all measurements after addition of oligomycin A was assumed to be zero. Finally, the ATP production rates *J*_ATP_total_, *J*_ATP_glycolysis_, *J*_ATP_oxphos_, *J*_ATP_TCA_, and *J*_ATP_mitochondria_ at measurement time *t* were calculated as follows:

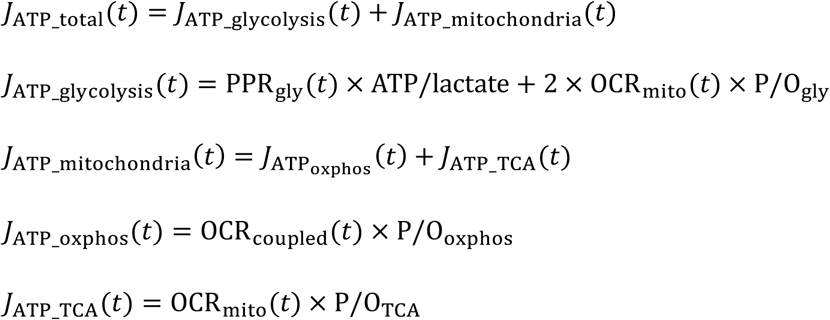

where ATP/lactate represents the ATP production per lactate produced. With glucose as the substrate, ATP/lactate is 1. With glycogen as the substrate, assuming a composition of 90% α-1,4-glycosidic bonds and 10% α-1,6-glycosidic bonds, the ATP/lactate is 1.45. The lactate production rate is assumed to be equal to PPR_gly_. The values of *max* H^+^/O_2_, ATP/lactate, P/O_gly_, P/O_oxphos_, and P/O_TCA_ for no substrate (i.e. glycogen), glucose and octanoate were taken from or calculated as in the works of Mookerjee et al. (Mookerjee et al., 2017, Mookerjee et al., 2015); these values are given in Supplementary Table 2.

#### Flux analysis of digitonin-permeabilized cells by Seahorse XF96

Cells were pre-incubated in a 96-well Seahorse culture plate in modified HBSS for ambient air with 5.5 mM glucose and 0.1% (w/v) fatty acid-free BSA (Suppl. Table 1) for 1 hour at 37°C in ambient air. Subsequently, 50 µM LRE1 or 0.1% DMSO (vehicle control) was added. After 10 minutes incubation, the media in all wells were replaced by MAS/BSA buffer (60 mM sucrose, 200 mM mannitol, 10 mM potassium phosphate buffer, 5 mM MgCl_2_, 1 mM EGTA, 20 mM HEPES and 0.4% fatty acid-free BSA, pH 7.1) as described by Salabei et al. (Salabei, Gibb et al., 2014) with the same substrates and inhibitors as in the previous incubation in HBSS. The plate was loaded onto the Seahorse XF96 Analyzer and the measurement started after a 10 minutes temperature equilibration. Cells were permeabilized with 25 µg/mL digitonin by injecting a concentrated mixture after baseline measurements. The injection mixture also contained substrates for mitochondrial respiration and 1 mM ADP (final concentration). The final concentrations of substrates are 5 mM succinate with 2.5 µM rotenone, 2 mM pyruvate with 1 mM malate, and 4 mM glutamate with 1 mM malate. When substrates were omitted, oxygen consumption decreased dramatically (data not shown). After about 15 minutes of ADP-driven state 3 respiration, oligomycin A (2.5 µM), FCCP (2.5 µM), and antimycin A (2.5 µM) plus rotenone (2.5 µM) were injected in order. The ADP-driven respiration rate was defined as the difference between the first OCR measurement after the addition the mixture of digitonin, ADP, and substrates and the last OCR measurement after the addition of oligomycin A. The FCCP-driven respiration rate was defined as the difference between the first OCR measurement after FCCP addition and the last OCR measurement after the addition of oligomycin A.

#### Statistics

All results are presented as mean ± standard deviation (SD). All *n* number represent the number of independent samples. Statistical significance was determined by two-tailed Student’s *t*-test, one-way analysis of variance (ANOVA) with Turkey’s or Dunnett’s multiple comparison test, or two-way ANOVA with Sidak’s multiple comparison test where applicable. Statistical analysis was performed with GraphPad Prism 7 (GraphPad Software, La Jolla, CA) with an α error of 0.05.

## Supplementary Figure Legends

**Figure S1.**
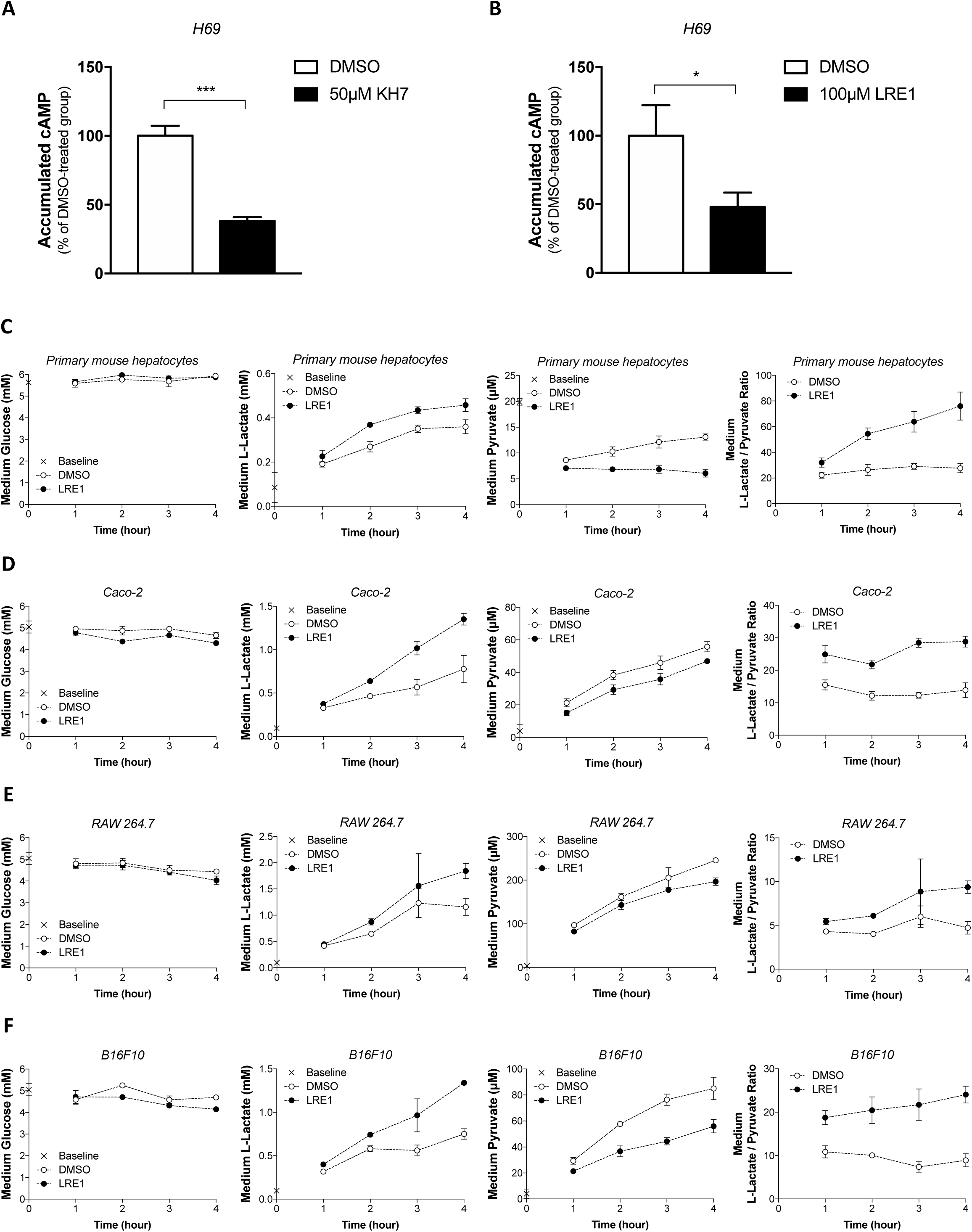
Suppression of sAC activity promotes glycolysis (related to Figure 1). (A-B) H69 cholangiocytes were treated with pan-phosphodiesterase inhibitor IBMX in the presence or absence of sAC inhibitor KH7 (A) or LRE1 (B) for 5 minutes. The accumulated cAMP was assayed by ELISA. Data represent mean ± SD (n = 3). (C-F) primary mouse hepatocytes (C), human colorectal carcinoma Caco-2 (D), Abelson murine leukemia virus-transformed macrophage Raw264.7 (E), and murine melanoma B16F10 (F) were treated with 50 µM LRE1 or 0.1% DMSO. Medium glucose, L-lactate, and pyruvate were sampled hourly for 4 hours and enzymatically determined. Data represent mean ± SD (n = 3). Statistical analysis: (A-B) Two-tailed unpaired Student’s *t*-test. *P<0.05, ***P<0.001.

**Figure S2.**
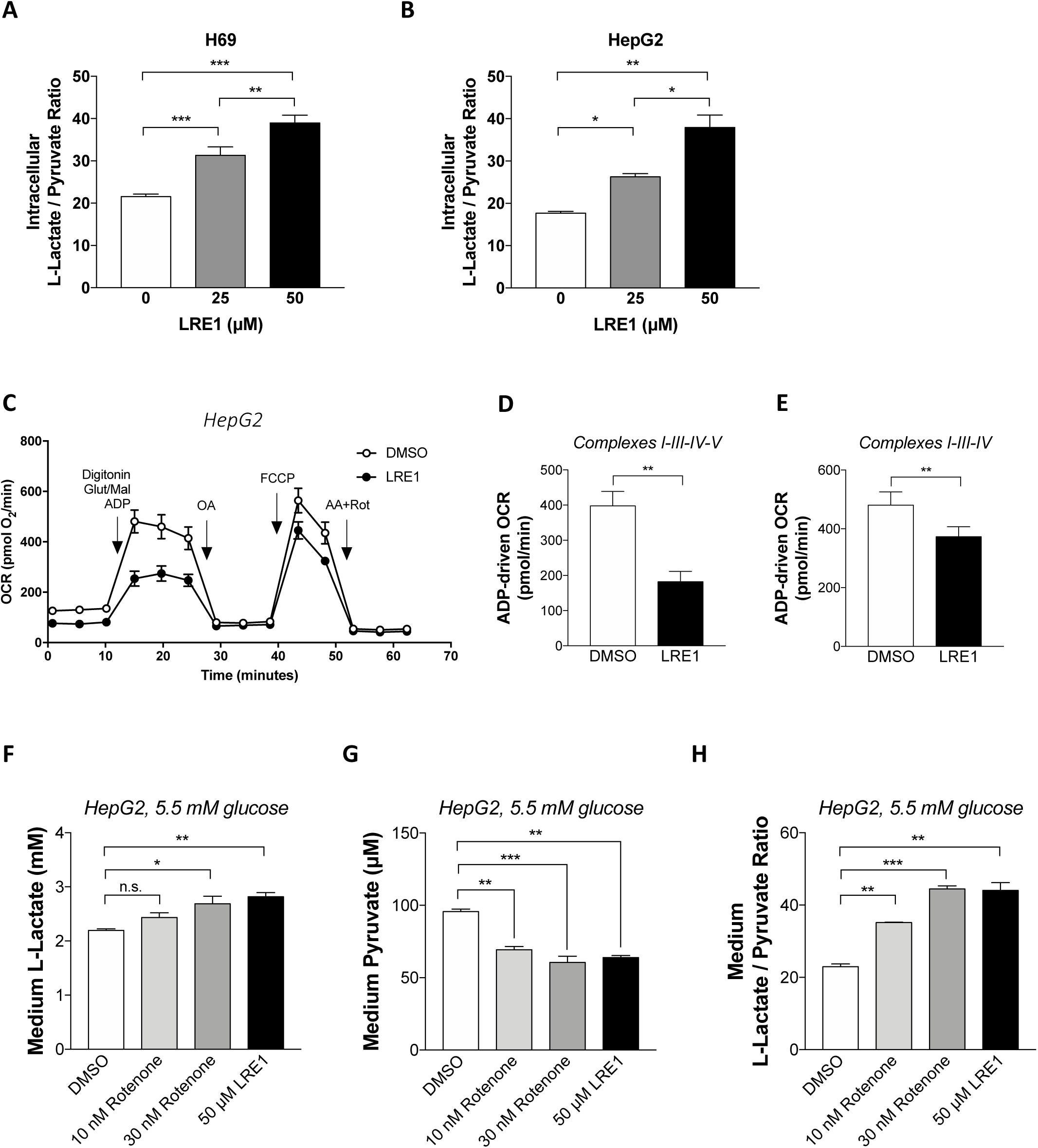
Soluble adenylyl cyclase regulates the cytosolic NADH/NAD^+^ redox state and aerobic glycolysis via complex I (related to Figure 2). (A-B) Intracellular lactate-to-pyruvate ratios in H69 cholangiocytes (A) and HepG2 cells (B) treated with 0.1% DMSO, 25 µM and 50 µM LRE1 for 6 hours. Data represent mean ± SD (n = 3 for A, n = 2 for B). (C-E) HepG2 cells were pre-treated with 0.1% DMSO or 50 µM LRE1 for 10 minutes in HBSS and then the medium was changed to MAS buffer. Oxygen consumption rate (OCR) was monitored by Seahorse Flux Analyzer XF96. Cells were permeabilized by adding 25 µg/mL digitonin together with 1 mM ADP, 4 mM glutamate, and 1 mM malate (n = 5 and 4 for DMSO and LRE1, respectively). Subsequently, 2.5 µM OA, 1 µM FCCP, and 2.5 µM antimycin A (AA) plus 2.5 µM rotenone (Rot) were injected as indicated. ADP-driven OCR (D) and FCCP-driven OCR (E) were derived from (C) as described in *Methods*. Data represent mean ± SD. (F-H) HepG2 cells were treated with 0.1% DMSO (vehicle control), 10 nM rotenone, 30 nM rotenone, and 50 µM LRE1 in medium containing 5.5 mM glucose for 1 hour. Medium L-lactate (F), pyruvate (G), and their ratio (H). Data represent mean ± SD (n = 2). Statistical analysis: (A-B) One-way ANOVA with Tukey’s multiple comparisons test. (D-E) Two-tailed unpaired Student’s *t*-test. (F-H) One-way ANOVA with Tukey’s multiple comparisons test. n.s., not significant; *P<0.05, **P<0.01, ***P<0.001.

**Figure S3.**
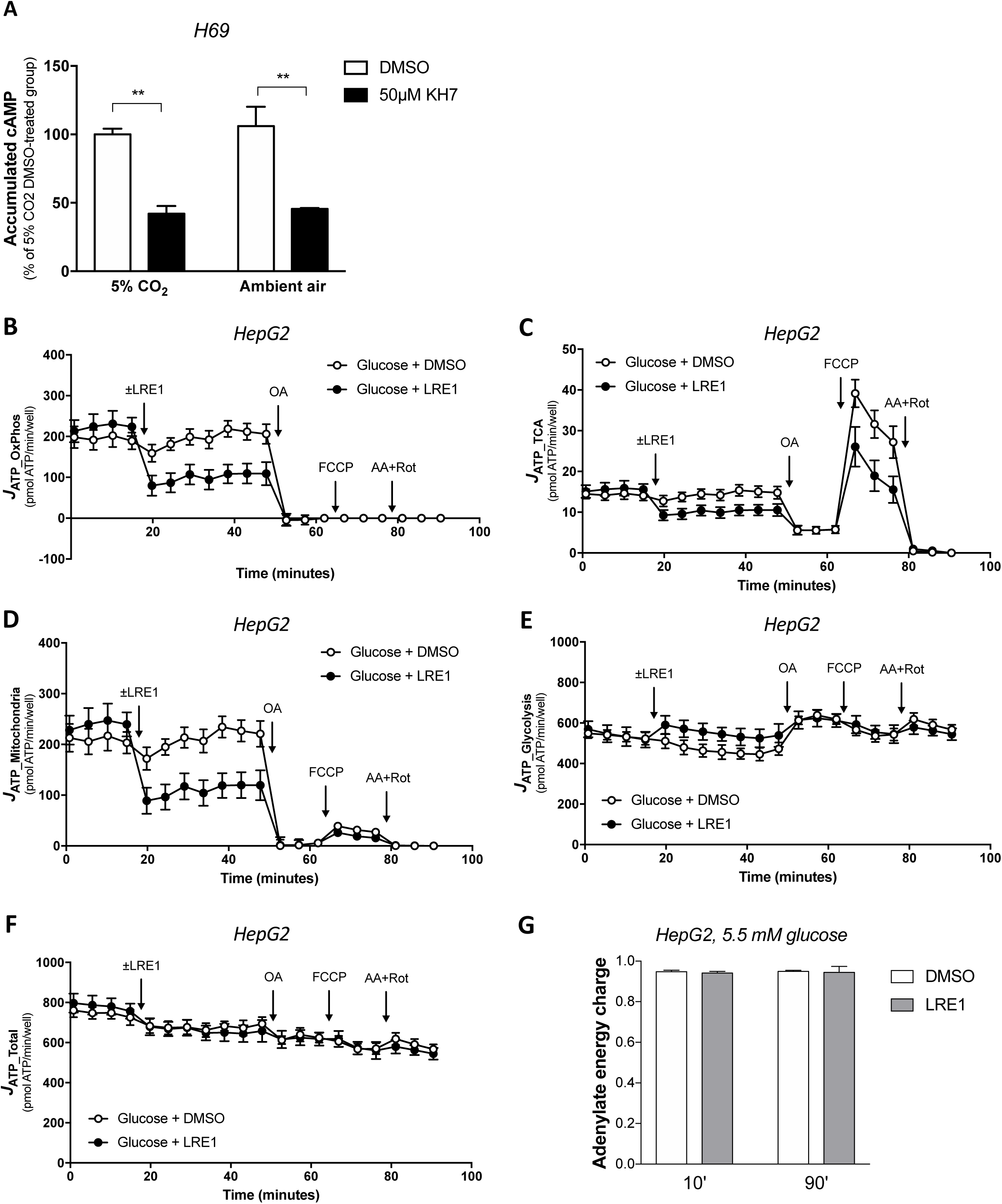
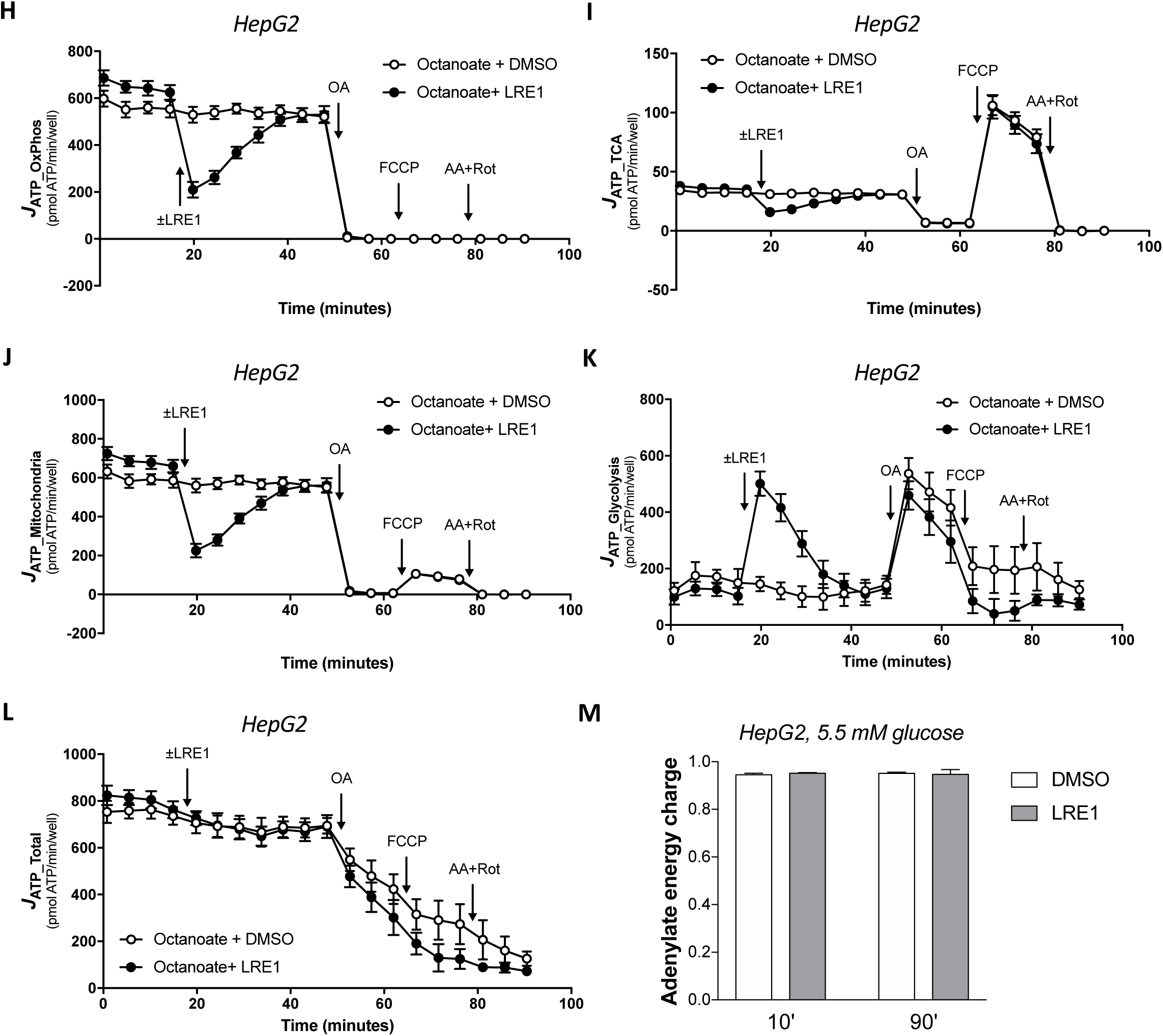
Soluble adenylyl cyclase is an acute switch for aerobic glycolysis that maintains energy homeostasis (related to Figure 3). (A) H69 cholangiocytes were pre-treated with sAC inhibitor (50 µM KH7) for 10 minutes and then pan-phosphodiesterase inhibitor IBMX was added to initiate cAMP accumulation in the presence or absence of in 5% CO_2_ incubator or in ambient air. After 6 minutes, the accumulated cAMP was assayed by competitive ELISA. Data is normalized to DMSO 5% CO_2_ group and represents mean ± SD (n = 2). (B-G) ATP production rates by oxidative phosphorylation (*J*_ATP_OxPhos_)(B), tricarboxylic cycle (*J*_ATP_TCA_)(C), mitochondria (*J*_ATP_Mitochondria_)(D), glycolysis (*J*_ATP_Glycolysis_)(E), and total ATP production rate (*J*_ATP_Total_)(F) were derived from OCR and ECAR measurements in Figure 2C and 2D as described in *Methods*. (G) HepG2 cells were treated with 0.1% DMSO (vehicle control) or 50 µM LRE1 for 10 and 90 minutes. AMP, ADP, and ATP were measured for the calculation of the adenylate energy charge. Data represent mean ± SD (n = 3). (H-L) ATP production rates by oxidative phosphorylation (*J*_ATP_OxPhos_)(H), tricarboxylic cycle (*J*_ATP_TCA_)(I), mitochondria (*J*_ATP_Mitochondria_)(J), glycolysis (*J*_ATP_Glycolysis_)(K), and Total ATP production rate (*J*_ATP_Total_)(L) were derived from OCR and ECAR measurements in Figure 3G and 3H as described in *Methods*. is defined as the sum of *J*_ATP_Glycolysis_ and *J*_ATP_Mitochondria_. (M) HepG2 cells were treated with 0.1% DMSO (vehicle control) or 50 µM LRE1 for 10 and 90 minutes. AMP, ADP, and ATP were measured for the calculation of the adenylate energy charge. Data represent mean ± SD (n = 3). Statistical analysis: (A) Two-way ANOVA with Sidak’s multiple comparisons test. *P<0.05.

**Supplementary Figure 4.**
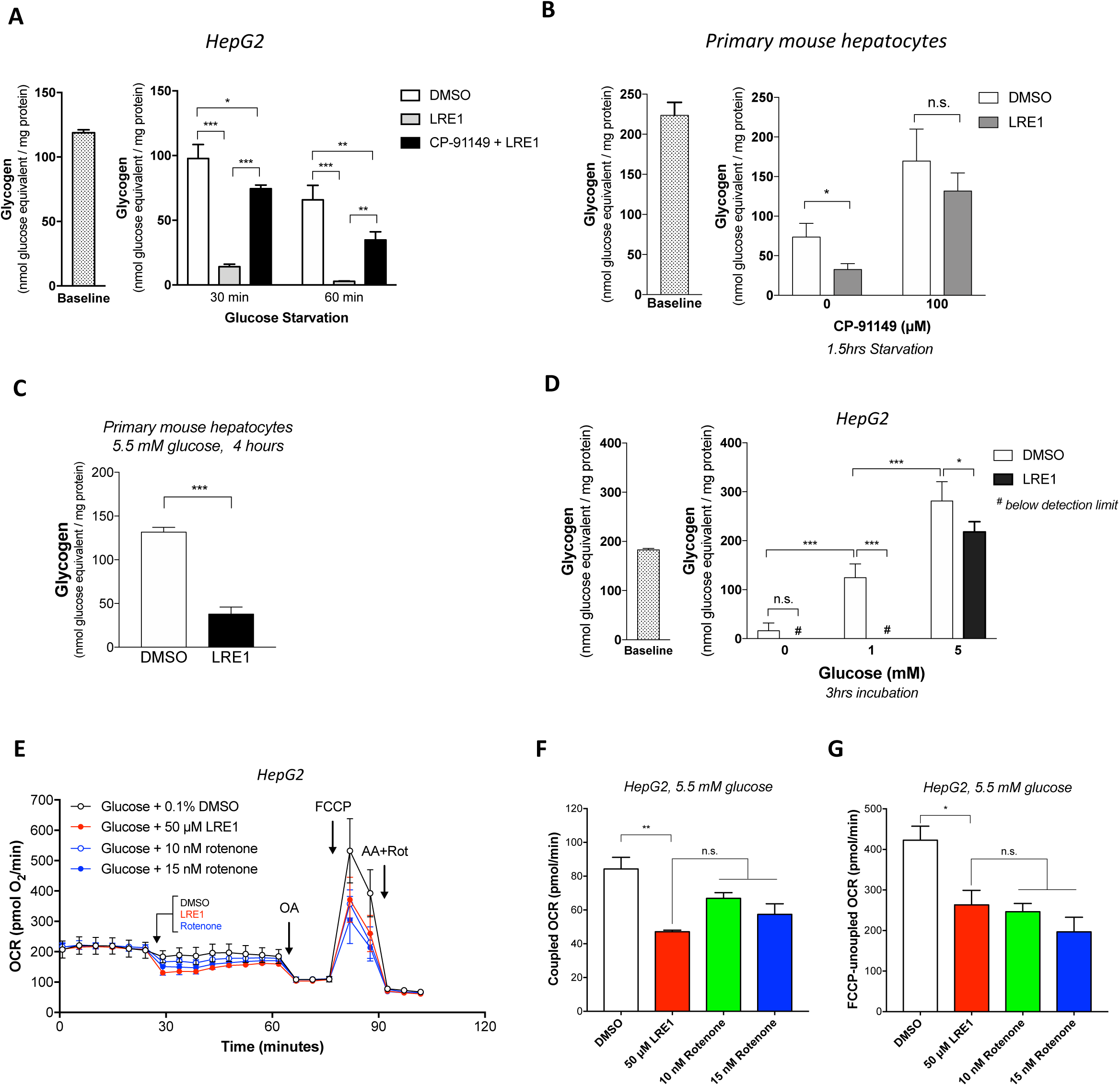
Inhibition of soluble adenylyl cyclase promotes glycogenolysis (related to Figure 4). (A) HepG2 cells acutely starved in the presence of 0.1% DMSO, 50 µM LRE1, or 50 µM LRE1 + 100 µM CP-91149. Glycogen contents were determined and normalized to protein content. Data are presented as mean ± SD (n = 3). (B) Primary mouse hepatocytes were acutely starved in the presence or absence of 50 µM LRE1 and 100 µM CP-91149 for 1.5 hours. Glycogen contents were determined and normalized to protein content. Data represent mean ± SD (n = 3). (C) Primary mouse hepatocytes were incubated with 50 µM LRE1 or vehicle control for 4 hours. Glycogen contents were determined and normalized to protein content. Data represent mean ± SD (n = 3). (D) HepG2 cells were incubated in medium with 0, 1, and 5 mM glucose in the presence or absence 50 µM LRE1 for 3 hours. Glycogen contents were determined and normalized to protein content. Data represent mean ± SD (n = 3). (E) HepG2 cells were pre-incubated for 60 minutes with HBSS with 5.5 mM glucose. Oxygen consumption rate was monitored. Arrows indicate the injection (in order) of 0.1% DMSO (n = 8), 50 µM LRE1 (n = 4), 10 nM rotenone (n = 4), or 15 nM rotenone (n = 5), oligomycin A (OA), carbonyl cyanide-*p*-trifluoromethoxyphenyl hydrazone (FCCP), and antimycin A (AA) and rotenone (Rot). The coupled respiration rate (F) and the FCCP-uncoupled respiration rate (G) were derived from (E) as described in *Materials and Methods*. # denotes values below the detection limit. Statistical analysis: (A-B) Two-way ANOVA with Sidak’s multiple comparisons test. (C) Two-tailed unpaired Student’s *t*-test. (D) Two-way ANOVA with Tukey’s multiple comparisons test. (F-G) One-way ANOVA with Dunnett’s multiple comparisons test (against DMSO group). n.s., not significant; *P<0.05, *P<0.01, ***P<0.001.

**Supplementary Figure 5.**
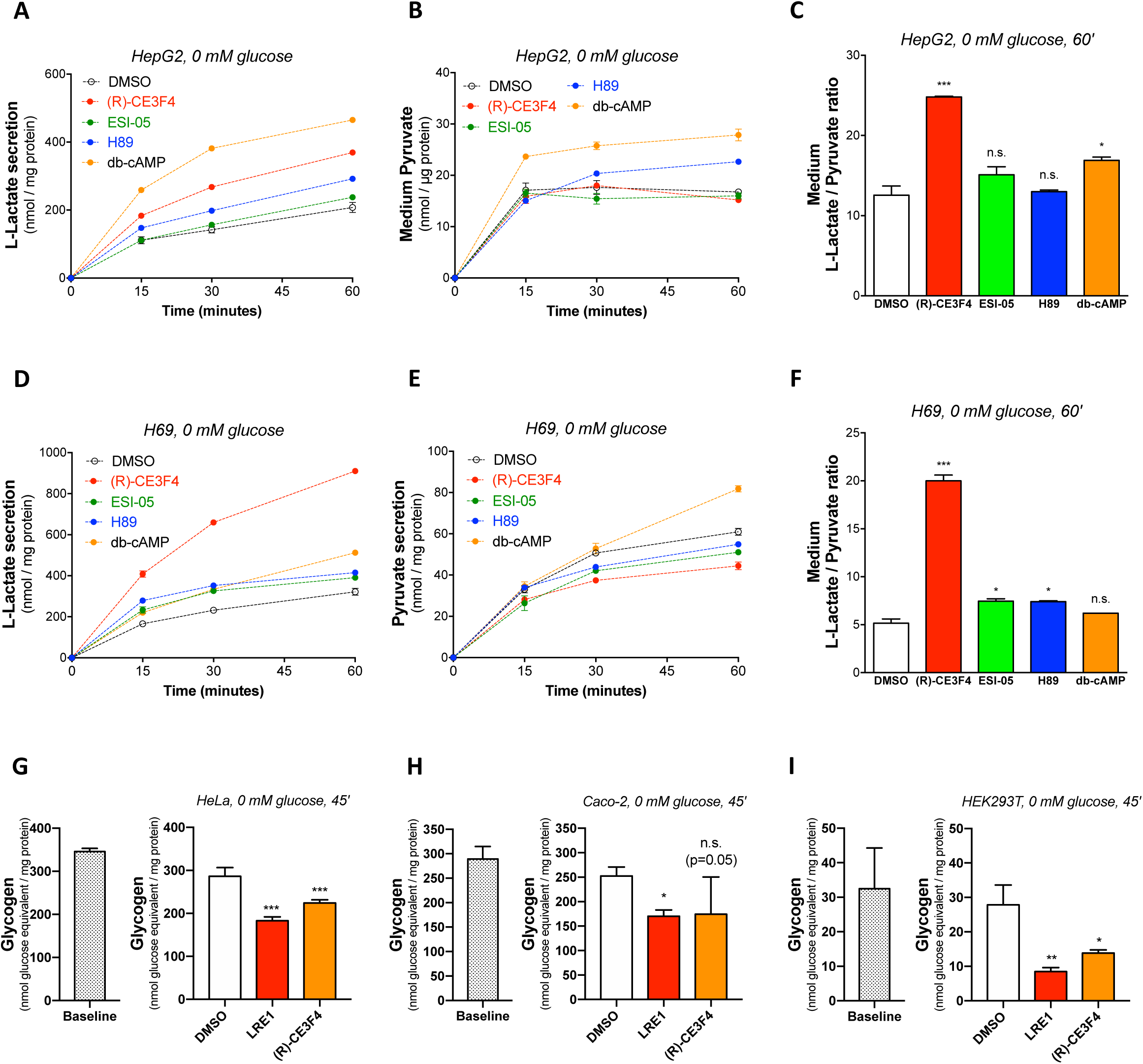
sAC-Epac1 and tmAC-PKA signaling have opposite effects on glycogen homeostasis (related to Figure 5). (A-C) HepG2 cells were acutely starved of glucose and treated with 0.1% DMSO (vehicle control), 50 µM (R)-CE3F4, 10 µM ESI-05, 10 µM H89, 100 µM db-cAMP. Medium L-lactate and pyruvate were assayed at indicated time points. Data represent mean ± SD (n = 2). (D-F) H69 cells were acutely starved of glucose and treated with 0.1% DMSO (vehicle control), 50 µM (R)-CE3F4, 10 µM ESI-05, 10 µM H89, 100 µM db-cAMP. Medium L-lactate and pyruvate were assayed at indicated time points. Data represent mean ± SD (n = 2). (G-I) HeLa cells (G), Caco-2 cells (H), HEK293T cells (I) were pre-incubated 15 minutes in glucose-free DMEM and then refreshed with glucose-free DMEM with 0.1% DMSO (vehicle control), 50 µM LRE1, and 50 µM (R)-CE3F4 for 45 minutes and then the glycogen content was determined. Data represent mean ± SD (n = 4). Statistical analysis: (C and F) One-way ANOVA with Dunnett’s multiple comparisons test (against DMSO group). n.s., not significant; *P<0.05, ***P<0.001. (G-I) One-way ANOVA with Dunnett’s multiple comparisons test (against DMSO group). n.s., not significant; *P<0.05, **P<0.01, ***P<0.001.

